# Inferences from epigenetic information in an ecological context: a case study of early-life environmental effects on DNA methylation in zebra finches

**DOI:** 10.1101/2025.07.30.667588

**Authors:** Marianthi Tangili, Per J. Palsbøll, Simon Verhulst

## Abstract

DNA methylation (DNAm) plays a key role in mediating phenotypic responses to environmental variation. Various approaches exist to link DNAm variation to phenotypes, ranging from single nucleotide resolution to the development of composite indexes (‘epigenetic clocks’). We here discuss the conceptual differences between these approaches through a case study using whole genome, longitudinal DNAm data from adult zebra finches raised in experimentally created small or large broods. Specifically, we (i) identified sex and age-specific CpG sites where DNAm was affected by brood size, and (ii) developed two DNAm-based indexes of the early-life environment (DMSi: based on differentially methylated sites with respect to rearing brood size; SMLmi: supervised machine learning-derived index optimized to predict brood size). We also compared results obtained by either merging or analyzing longitudinal DNAm samples separately and discussed how the permanence or transience of DNAm changes shapes responses to environmental variation through life. Our findings confirm that early-life environment leaves lasting DNAm signatures detectible in adulthood, and this effect is stronger later in life. Importantly, both indexes predicted early-life growth rates, demonstrating that DNAm-based indexes can be leveraged to retroactively quantify aspects of early-life conditions, providing a powerful novel tool in the study of wild populations.

## 1. Introduction

Evolutionary theory has long considered genetic variation to be the main driver of phenotypic plasticity. However, mounting evidence reveals the insufficiency of genetic variation in explaining the rapid phenotypic responses of wild animals to environmental changes (Charmantier et al., 2008; Réale et al., 2003). Epigenetic modifications are known to be sensitive to the environment (Angers et al., 2010; Li et al., 2021), modulate gene expression (Gibney & Nolan, 2010), and generate phenotypic variation without altering the primary nucleotide sequence with direct effects on evolution and adaptation (Kronholm & Collins, 2016). Emerging research also recognizes epigenetic mechanisms such as chromatin modifications and DNA methylation (DNAm) as possible regulators that modulate phenotypic plasticity and thereby link environmental variation to phenotypic outcomes. Epigenetic modifications are especially relevant in contexts when genetic variation is limited, potentially generating heritable phenotypic diversity that can be subject to natural selection (Lind & Spagopoulou, 2018).

Since methylation of different parts of the genome has different effects on gene expression (Moore et al., 2013), τhe design and resolution of DNAm studies can greatly impact the insights that can be drawn. All levels of analysis complement each other, providing a comprehensive understanding of the DNAm landscape and how that influences gene expression, cellular function and ultimately individual phenotype across different biological contexts. Global DNAm, which provides a single overall measure of DNAm across the entire genome of an individual (e.g. with gel-based techniques or ELISA kits) is a simple, fast and cost-effective way of studying DNAm. It can provide a broad overview of the epigenetic state of individuals, cells or tissues, which can be indicative of overall genomic stability and health (reviewed by Li & Tollefsbol, 2021). However, effects of age and environmental factors can result in increased DNAm at some CpG sites, while simultaneously resulting in DNAm decreases at other sites. As a consequence, whole genome DNAm levels are much coarser and less informative as they are skewed by repetitive regions and cannot be directly connected to regulatory pathways as evidenced by previous research in mammals and birds (Laubach et al., 2021; Meyer et al., 2023).

Higher-resolution approaches assess DNAm at the level of individual CpG sites (positions in the genome where a 5’ cytosine is followed by a guanine, Moore et al., 2013) providing valuable insights, especially when directly linked to gene expression of a gene with known effects on the phenotype or life history of the organism. In both biomedical and ecological contexts, DNAm patterns of specific CpG sites have been associated with numerous traits. In clinical research, CpG-specific DNAm patterns have been linked to specific diseases (Hedman et al., 2017) and neuropsychiatric disorders (Zheleznyakova et al., 2016). In evolutionary ecology, the methylation of specific CpG sites, or entire genes has been studied in relation to a plethora of ecologically relevant traits across a wide variety of species. In birds, methylation has been associated with brood size of rearing (Jimeno et al., 2019; Sepers et al., 2021), food deprivation (Xiao et al., 2020), and urbanization (Caizergues et al., 2022). In wild baboons, methylation was linked to resource availability (Lea et al., 2016), in lizards to habitat quality (Hu et al., 2019), and in crustaceans to the exposure to pollution (Harney et al., 2022). Collectively, these findings confirm that DNAm is sensitive to environmental perturbations and a putative mechanism for generating phenotypic variation.

A fundamentally different approach to make inferences based on DNAm, compared to the focus on individual CpG sites, is the development of methylation-based indexes. Utilizing the methylation of selected positions in the genome, an algorithm can be trained and later used as a predictive tool using e.g. supervised machine learning. Such indexes have proven to be valuable for diagnostic and therapeutic purposes, for example for non-invasive cancer detection and identifying cancer types (Draškovič & Hauptman, 2024; Wei et al., 2021), prediction of patient response to chemotherapy (Wick et al., 2014), and assessment of the impact of lifestyle and therapeutic interventions (Keller et al., 2020; Lu et al., 2023). This approach has in particular been applied to develop epigenetic ‘clocks’ of chronological age, algorithms that use DNAm patterns to predict chronological age for humans, model, and non-model animals (Horvath, 2013; Petkovich et al., 2017; Tangili et al., 2023).

Despite the growing use of DNAm based indexes, it is still debated whether the CpG sites underlying such indexes are functionally involved in the targeted process, or if the level of methylation at the identified sites correlates with those processes in absence of any biological function. The functional role of the DNAm of CpG sites outside of regulatory regions remains poorly understood (Moore et al., 2013) and confounded by the limited overlap between the CpG sites identified across different studies aimed at the same biological process within the same species (Galkin et al., 2020; Liu et al., 2020). Part of the explanation might be that phenotypic variation rarely correlates strongly with single genes and hence methylation-based indexes likely capture CpG sites across gene networks scattered across the genome. In this sense, an index comprising DNAm at many CpG sites may better match the causes of the phenotypic variation under study, compared to the DNAm of single sites. Nevertheless, whether these sites are causally linked to the observed phenotypes remains an open question, demanding more research across diverse species to elucidate the underlying mechanisms.

We here present a case study to explore different approaches to utilize DNAm data to gain information on the role of DNAm in mediating the effects of the environment on the phenotype. We have utilized longitudinal whole genome methylation data from fifty adult zebra finches (*Taeniopygia castanotis*) of both sexes that were reared in experimentally created small or large broods. This manipulation is of interest, firstly because it is reasonable to assume that parents and offspring have evolved responses to variation in brood size because it shows strong natural variation, and hence we can reasonably assume that phenotypic effects are not artefacts. Secondly, previous studies from the present study population and others, have documented multiple long-term effects of the nutritional stress caused by this manipulation; for example, individuals reared in large broods have shorter lifespans (Briga et al., 2017) and lower sexual attractiveness (Holveck & Riebel, 2010).

Longitudinal data, collected from the same individuals at multiple time points, are essential when studying processes such as development, aging and adaptation and allow for the continuous monitoring of the molecular state of an individual. However, such data is scarce and most epigenetic studies are cross-sectional. Knowing how to analyze and interpret longitudinal epigenetic data is crucial for understanding complex phenotypic traits, but there is as still little experience with utilizing longitudinal samples in epigenetic studies. In our case study we sampled all individuals twice, once in early adulthood, and a second time closer to the end of their lives, and explored different ways to combine the information of longitudinal samples.

In our case study, we first identified CpG sites with differential methylation between treatments, for the sexes pooled but also for the sexes separately, to identify sex-specific effects of developmental conditions. Secondly, to translate methylation information into interpretable metrics, we developed two methylation-based indexes: an epigenetic early life quality index (DMSi) calculated from methylation data of differentially methylated sites between individuals raised in small and large broods and a predicted brood size index (SMLmi) calculated using a supervised machine learning algorithm with the goal to predict brood size of rearing. We hypothesized that these indexes could serve to quantify effects of early life conditions on the methylome within the treatment categories, and as such provide a more accurate reflection of an individual’s phenotypic state than the coarse binary trait of manipulated brood size. If successful, an estimate could in this way be obtained of the early life conditions experienced by individuals first encountered in adulthood, providing a new tool for studies in the wild where a substantial part of the population can consist of immigrants with unknown history.

## 2. Case study

### 2.1 Case study: methods

#### 2.1.1 Subjects and samples

The present data set is based on samples taken from birds in the hard foraging conditions in the experiment reported in (Briga et al., 2017), where methods are described in detail. In brief, adult zebra finches were randomly matched and provided with nest boxes and nesting material. Fortified canary food was supplied to the parents in addition to tropical seed mixture and water until the hatching of the first chick in the nest but not thereafter. Nestboxes were checked daily for the presence of eggs or chicks. Chicks were cross-fostered at age five days or before to broods of either two or six chicks, which is the range of brood sizes observed in the wild (Zann, 1996). On the day of cross-fostering, all chicks were weighed and individually marked, and weighed again when 15 days old (or as near to that as practically possible). Siblings were assigned to different broods, and parents never reared their own offspring. When reaching adulthood (≥3 months), birds were moved to outdoor aviaries each containing single sex groups of 18–24 adults. We here used blood samples collected from the brachial vein and stored in -80^°^C in glycerol buffer from 50 individuals distributed equally over the small and large broods (n=25 x 2) and almost equally over the two sexes (26 males, 24 females). Given all other selection criteria, we could not make within natal family comparisons with respect to the effect of rearing brood size, although this would have further increased statistical power. Each bird was sampled every six months after entering the experiment, and DNAm data were extracted for two samples per individual collected early and late in their life, with an average sampling interval of 995 (S.D.=742) days - see Fig.S1 for the age distribution at sampling.

#### 2.1.2 Enzymatic Methyl-seq data

We extracted DNA according to the manufacturer’s protocol using innuPREP™ DNA Mini Kit (Analytik Jena GmBH) from 3 uL of nucleated red blood cells. Next-generation sequencing was conducted by the Hospital for Sick Children (Toronto, Canada). The amount of DNA was quantified using a Qubit High Sensitivity Assay (Thermo Fisher Scientific). Two hundred ng of DNA was used as input material for library preparation using the Next Enzymatic Methyl-seq™ Kit (New England Biolabs Inc.) following the manufacturer’s instructions. Briefly, DNA was fragmented by sonication to an average length of ∼500 base pairs (bp) using a Covaris LE220 (Covaris Inc.). Fragmented DNA was end-repaired and ligated to Illumina sequencing adapters. The 5-methylcytosines and 5-hydroxymethylcytosines were oxidized by TET2 and cytosines were deaminated by APOBEC. Methyl-seq libraries were subjected to five PCR amplification cycles. Libraries assessed on a Bioanalyzer™ DNA High Sensitivity chip (Agilent Inc.). The amount of DNA was quantified by quantitative PCR using the Kapa Library Quantification Illumina/ABI Prism™ kit according to the manufacturer’s instructions (La Roche Ltd.). Libraries were pooled in equimolar amounts and sequenced to a minimum of ∼100M paired-end reads (2×150bp) per sample on a 10B flow cell using a NovaSeqX™ (Illumina Inc.) platform.

Sequences were trimmed using Trim Galore! v0.6.10 (Krueger et al., 2023) in paired-end mode. The data were assessed before and after trimming using FastQC v. 011.9 (Andrews, 2010) and MultiQC v. 11.14 (Ewels et al., 2016). Trimmed reads were aligned against the *in silico* bisulfite converted zebra finch (*Taeniopygia castanotis*) reference genome (GCA_003957565.4, Rhie et al., 2021) using Bismark v. 0.14.433 (Krueger & Andrews, 2011) using the Bowtie 2 v. 2.4.5 alignment algorithm (Langmead & Salzberg, 2012). The average mapping efficiency was at 65.14% (SD: 2.79). Mean autosomal DNAm did not appear to be affected by the storage time (range 8-16 years) of the samples (estimate= 0.03 ± 0.04, t = 0.85, *p* = 0.39).

#### 2.1.3 Differentially methylated site (DMS) identification

We performed multiple epigenome wide association studies (EWASs) in order to identify differentially methylated sites (DMSs) between zebra finches raised in small and large broods using *Methylkit* v. 1.18.0 (Akalin et al., 2015).

For all EWAS analyses the following steps were followed:

1. CpG sites were filtered for coverage, retaining those with a minimum coverage of 5 and maximum coverage the 99.9^th^ percentile coverage in order to avoid PCR bias (Wreczycka et al., 2017). For the analysis where the two samples from the same individual were merged, a minimum coverage filter of 12 was applied.
2. Read coverage distributions across all samples were normalized using the *normalizeCoverage* function of the *Methylkit* package.
3. Only CpG sites shared by all samples were retained using the *unite* function in the *Methylkit* package.
4. Invariable CpG sites with a methylation of zero or one hundred percent in all samples were excluded.
5. Logistic regression models with overdispersion correction following the McCullagh and Nelder method (“MN”, McCullagh & Nelder, 1989) were used to test for significant methylation differences between birds raised in small and large broods, accounting for extra-binomial variation in read counts. Resulting *p*-values were adjusted for multiple testing using a Bonferroni correction (Dunn, 1961).

We considered as DMSs all sites with a significant (≤0.05) Bonferroni-corrected *p*-value.

In order to explore the long-term effects of brood size on DNAm per CpG site, data of the two observations per individual were merged by adding the amount of methylated counts and dividing by the combined coverage of the two observations. Merging the DNAm data per individual increases coverage and therefore presumably precision.

When analyzing individual samples, filtering retained 1,114,704 CpG sites, whereas merging two samples per individual increased the number to 2,085,308 sites. Running the analysis by sex resulted in 1,896,932 and 1,787,898 CpG sites retained for females and males, respectively. After filtering, 60,046 CpG sites remained in the analysis for the first sample of all individuals (young age), and 82,273 sites in the second sample of all individuals (old age).

To assess whether age at sampling of the zebra finches influenced the differential methylation calculation, we adding age as a covariate (in all analyses using *Methylkit* except from the one where samples from the same individual were merged). The inclusion of age at collection as a covariate did not affect the results (methylation difference, significance) so it was omitted from subsequent analyses.

#### 2.1.4 Epigenetic early-life quality index (DMSi) calculation

We calculated an epigenetic early life quality index, which we termed DMSi, as follows: DNAm of all identified DMSs for the analysis per sample as mean centered, and for sites where DNAm was on average higher in birds from enlarged broods, mean centered DNAm was multiplied by -1. DNAm was then averaged per sample or individual over all DMSs. To facilitate comparisons with future studies we z-scored the index prior to analyses.

#### 2.1.5 Supervised Machine Learning methylation index (SMLmi) calculation

As a second method to develop a DNAm index of early life quality, we performed an elastic net regression using DNAm at variant sites (i.e. DNAm not of zero or one hundred in all samples) with a coverage between ten and the 99.9^th^ percentile of coverage. We used DNAm per CpG site per sample as the predictor variable and brood size of rearing as continuous dependent variable (coded as -0.5 and +0.5 for small and large rearing brood size respectively). The elastic net mixing parameter was set to α = 0.1, as a titration procedure showed this value to yield the most accurate model (i.e. most samples assigned to the correct brood size). To determine the optimal regularization parameter λ, we first used the function *cv.glmnet* from package *glmnet* v. 4.1-8 (Friedman et al., 2010) with a 10-fold cross-validation. The selected λ was the applied in a leave-one-out cross-validation (LOOCV) framework, in which one sample was omitted each time and the model was re-fitted on the remaining data, and the brood size of the omitted sample was predicted. Lastly, a final elastic net model was fit using all samples and the optimized hyperparameters. CpG sites with non-zero regression coefficients were extracted as features retained by the final model.

#### 2.1.6 Statistical analyses

We implemented linear mixed effects models using package *lme4* v. 1.1.35.1 (Bates et al., 2015) in R v. 4.1.2 (R Core Team, 2023) in order to explore how the DMSi or SMLmi differed between the sexes and individuals raised in small and large broods. We employed Akaike’s An Information Criterion (AIC) values (Akaike, 1974) to compare the predictive performance of models. Partial correlations were calculated using the *pcor* function from the *ppcor* v. 1.1 package (Kim, 2015) in R.

### 2.2 Case study: differentially methylated sites (DMSs)

When analyzing longitudinal DNAm data to extract differential methylation information, several analytical strategies can be applied. We identified DMSs between birds raised in small and large broods (i) separately for the two samples from each individual, (ii) merged per individual, (iii) separately for the first and second samples per individual (young and old) and (iv) separately for males and females. In this way, we can evaluate the effect of pooling samples taken at different ages on the detection of DMSs, on the consistency of DMSs over life, and on the age-dependence of frequency and magnitude of DMSs. Moreover, we can get insights into sex-specific effects of developmental conditions on DNAm.

#### 2.2.1 DMSs per sample vs. individual

When considering individual samples, the range of percent methylation difference between zebra finches raised in small and large broods was from -34.1 to 34.1% (Fig.1a). We identified 27 DMSs (Fig.1a), 20 of which were located in chromosome 3, two in chromosome 1A, two in chromosome 2 and one each in chromosomes 1, 10 and 7.

We extended the above analysis by combining the methylation data from the two samples collected early and late in life for each individual. The range of percent methylation difference between zebra finches raised in small and large broods was -34.5 to 34.2% (Fig.1b). However, this approach did not yield any DMSs between individuals raised in small and large broods (Fig.1b).

When comparing the absolute difference in methylation of all CpG sites in the analysis per sample vs. the merged samples using a Wilcoxon rank-sum test, this revealed a small but highly significant difference between groups (Fig.S2; W = 1.15 × 10^12^, *p*<0.001), with higher absolute differences between birds raised in small and large broods observed in the merged dataset. This is somewhat paradoxical, given that there are fewer DMSs identified in the pooled samples, for which we have no clear explanation. Importantly, these results indicate that effect-sizes are sensitive to how repeated measurements are treated, with the apparent gain in precision due to a higher coverage potentially outweighed by changes over life overshadowed by pooling samples.

#### 2.2.2 DMSs in young vs. old cohorts

The range of percent methylation difference between zebra finches raised in small and large broods was from -19.1 and 21.1% between young (Fig. 2a) and -25.8 and 28.41% between old birds (Fig.2b). After filtering, 49,538 sites (of the 60,046 in the young and 82,273 in the old cohort) were shared between the young and old cohort data sets, Across all 49,538 CpG sites, methylation differences between zebra finches raised in small and large broods were positively among age cohorts (Fig.S3, r= 0.29, *p* < 0.001), indicating a significant repeatability over life of the brood size effect on DNAm. However, we identified zero brood size-related DMSs when individuals were respectively young or old (Fig.2). CpG sites with the largest methylation differences between small and large broods were not shared by the young and old datasets. In samples taken early in life, there were more methylation differences around zero, while the most extreme methylation differences are found in the old cohort (Fig.S4). Indeed, when comparing the absolute methylation difference over all CpG sites using a Wilcoxon rank-sum test, this revealed a small but highly significant difference between samples taken early and late in life (Fig.S5; W = 2.16 × 10^9^, *p* < 0.001). Thus, the magnitude of the effect of early-life conditions on DNAm differed significantly within individuals over life, with the changes being more pronounced later in life, but, at the same time, no significant DMSs were observed when considering samples taken early and late in life separately.

**Figure 1.**
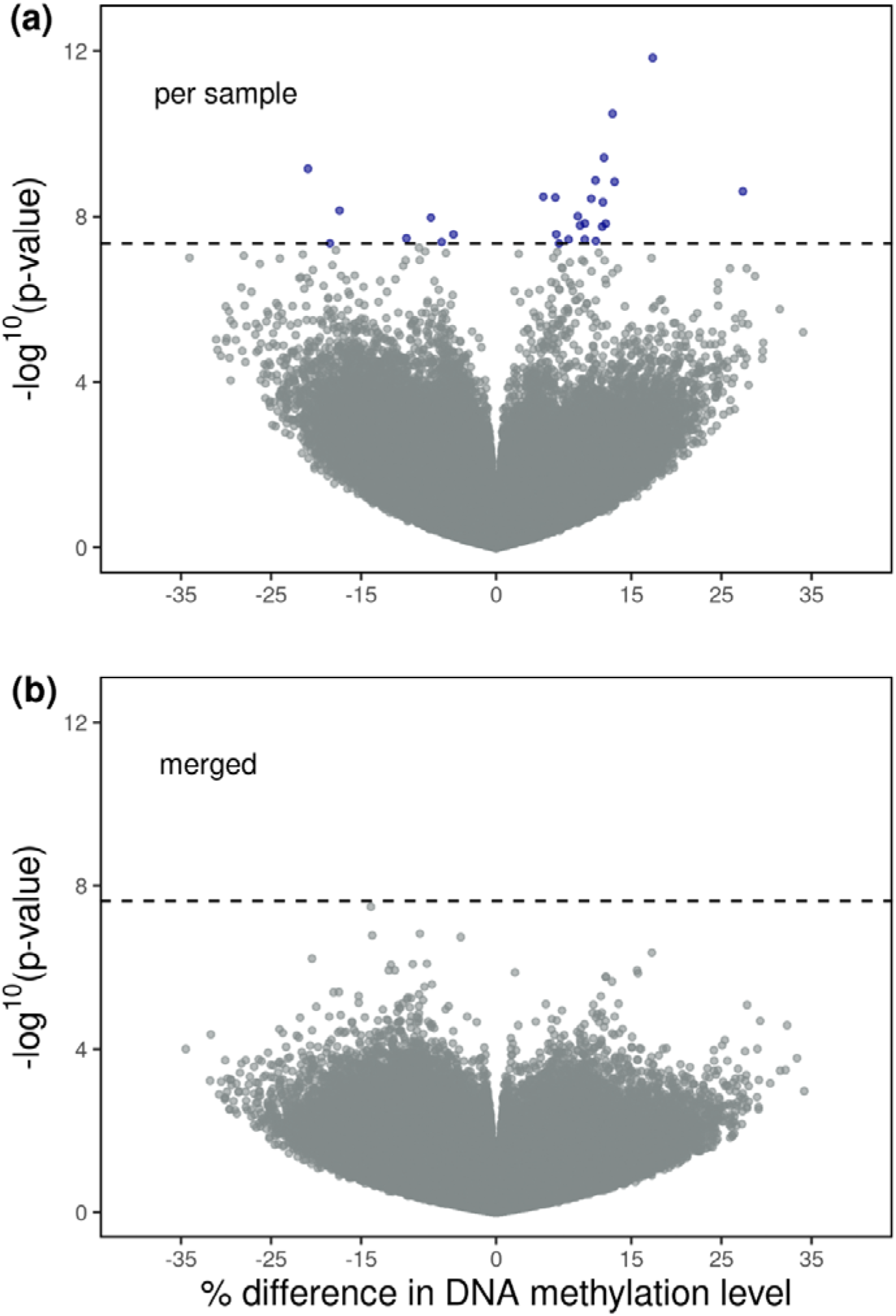
Volcano plot of Bonferroni-corrected p-values as a function of % DNAm differences between individuals raised in small and large broods for (a) the analysis performed per sample (N=1,114,704) and (b) for individuals merged together (N=2,085,308). Each point represents a CpG site, with colored points indicating statistically significant differences in % DNAm between individuals raised in small and large broods. The dashed horizontal lines mark the genome wide significance thresholds.

**Figure 2.**
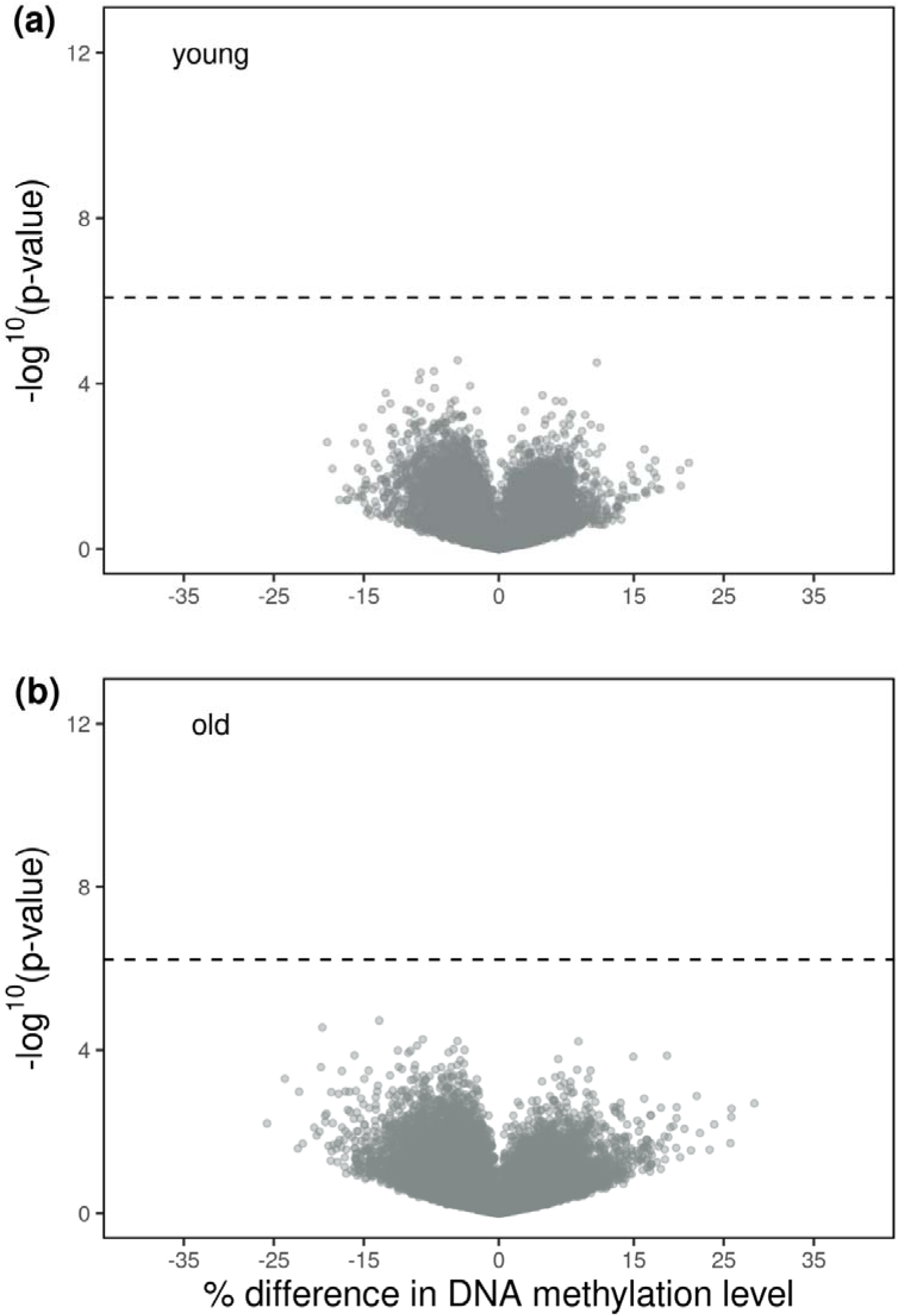
Volcano plot of Bonferroni-corrected p-values as a function of % DNAm differences between individuals raised in small and large broods for (a) the analysis performed only for the young cohort (N= 60,046) and (b) only for the old cohort (N=82,273). Each point represents a CpG site. The dashed horizontal lines mark the genome wide significance thresholds. For information about the age distribution of samples in each age cohort see Fig.S1.

#### 2.2.3 DMSs in females vs. males

The analysis for identification of DMSs was run separately for the two sexes. The range of percent methylation difference between zebra finches raised in small and large broods was from -48.6 and 47.4% in females (Fig.3a) and from -43.5 to 45.3% for males (Fig.3b), i.e. considerably larger then when observed when the sexes were pooled (section 2.2.1). After filtering, 1,108,263 CpG sites were shared between males and females. Across all 1,108,263 CpG sites, there was a weak but significant negative correlation between brood size effects on DNAm in females and males (Fig.S6, r = -0.04, *p*<0.001).

We identified 34 DMSs between females and 38 DMSs between males raised in small and large broods (Fig.3). The majority of female DMSs (N=16) were located in chromosome 4 while the majority of male DMSs (N=19) were located in chromosome 3. Moreover, in females, the majority of DMSs exhibited negative methylation differences (7 positive vs. 27 negative) whereas in males, the majority were positive (30 positive vs. 8 negative). This difference in distribution was highly significant (χ² = 22.18, df = 1, *p* < 0.001). These findings indicate a sex-specific pattern in methylation changes between zebra finches raised in small and large broods.

**Figure 3.**
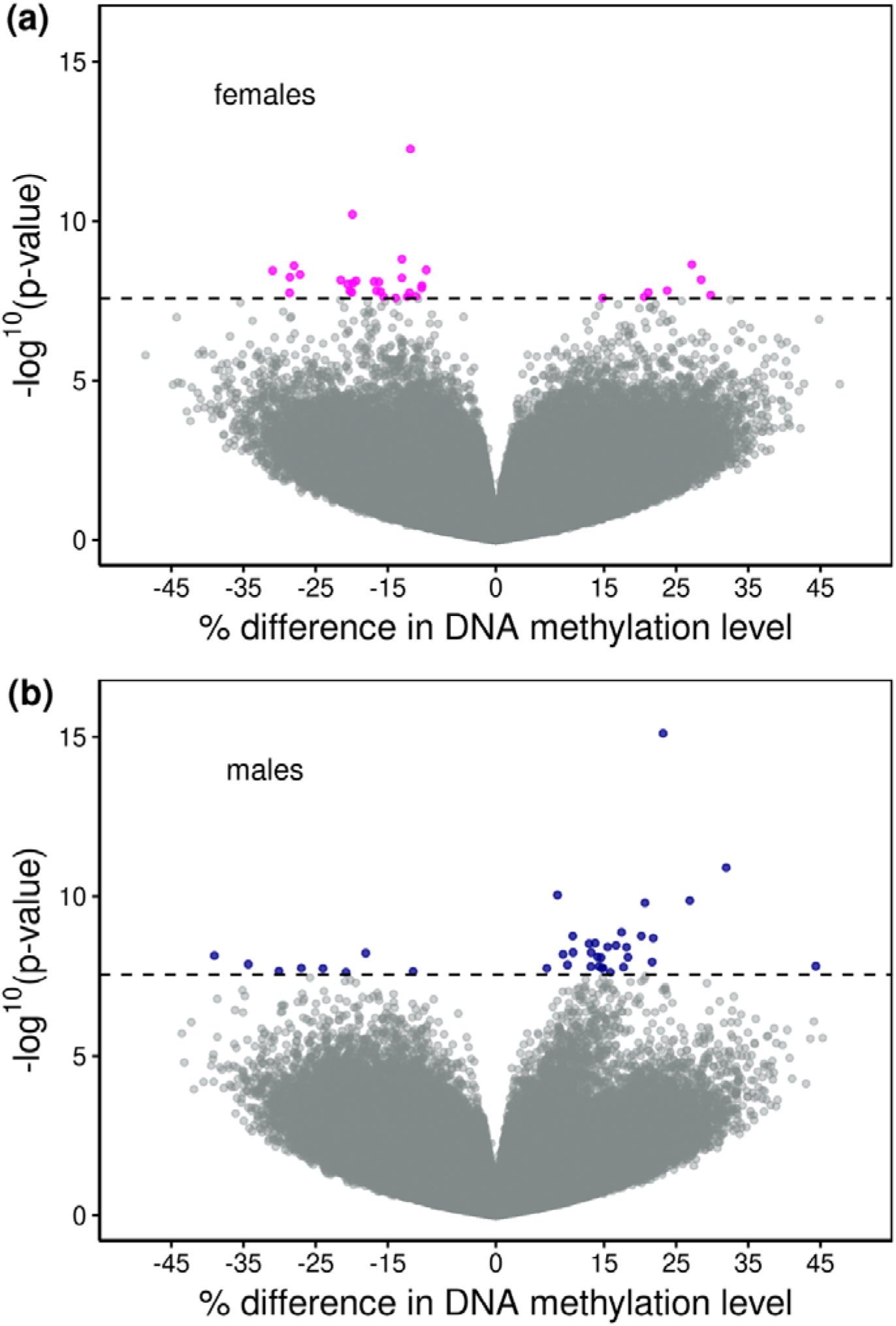
Volcano plot of Bonferroni-corrected p-values as a function of % DNAm differences between individuals raised in small and large broods for (a) the analysis performed only for females (N= 1,896,932) and (b) only for males (N=1,787,898). Each point represents a CpG site, with colored points indicating statistically significant differences in % DNAm between individuals raised in small and large broods.

In addition to the sex difference in average direction of DMSs, the absolute effect of brood size on all CpG sites was also sex-dependent. When comparing the absolute difference in methylation between birds raised in small and large broods across all CpG sites that passed the filters for females (N = 1,896,932) and males (N = 1,787,898) using a Wilcoxon rank-sum test, this revealed a small but highly significant difference between groups (Fig.S7; W = 1.64 × 10^12^, *p*<0.001), with females having larger absolute differences in methylation (Fig.S7). The above observations indicate sex-specific epigenetic responses to early-life environment.

The identification of CpG sites across the genome with significant methylation differences between adult zebra finches raised in small and large indicates that early-life conditions are associated with persistent, site-specific variation in DNAm across the genome. Previous work on nestling zebra finches has also shown consistent effects of brood size on DNAm (Jimeno et al., 2019; Sheldon et al., 2018). These changes in DNAm between individuals raised in different brood sizes might underlie the phenotypic differences observed between treatment groups such as the significantly shorter lifespan of zebra finches raised in large broods we have previously identified (Briga et al., 2017). Using longitudinal samples, that are therefore immune to effects of selective disappearance, we further extend these findings, by revealing that early-life environment effects on DNAm are stronger later in life. This raises the question whether early-life environmental effects on the phenotypic level also increased with age, but this remains to be investigated. In any case, our findings highlight the potential role of DNAm in mediating the enduring impact of early-life environmental conditions on adult phenotypes (Dettmer & Chusyd, 2023).

### 2.3 Case study: Differentially Methylated Site index (DMSi)

Methylation-based, predictive indexes are widely used in clinical research and diagnostics (Xu et al., 2024; Yousefi et al., 2022; Zhang et al., 2025), but have so far only been used to predict chronological age in an ecological context (Tangili et al., 2023). Developing methylation-based biomarkers could allow quantification of other traits that are difficult to estimate directly in individuals not previously encountered, e.g. early-life stress and other significant life events. The development of such biomarkers holds promise especially when studying wild populations, where detailed, individual life-history information might be labor-intensive to obtain and therefore scarce. The development of targeted, cost-effective methods to measure DNAm of selected CpG sites (e.g. Morselli et al., 2021), makes feasible the collection of population data with unprecedented accuracy.

We constructed an epigenetic index of early developmental conditions (i.e. brood size) using the methylation of the DMSs identified between zebra finches raised in small and large broods (DMSi) using individual samples (N=27). Because birds raised in large broods achieved a shorter lifespan (Briga et al 2017), we hypothesized that this index would serve as a quantitative measure of the overall phenotypic quality of each individual.

DMSi calculated over samples collected early and late in life on the same individuals was highly correlated (Fig 4a, r = 0.99, *p* < 0.001), indicating consistence of the developmental effects on DNAm over life. The DMSi differed strongly between birds reared in small and large broods, as expected (Fig. 4b-d). Interestingly, there was also a large difference in the DMSi variance between birds reared in small (0.04) and large (1.08) broods (F = 0.038, 95% C.I. 0.021 - 0.066, d.f. = 49, *p* < 0.001). We infer this to reflect that development in small broods is less variable compared to large broods, due to nutritional stress being more likely to occur in large broods, with larger variation between individuals in how they are impacted due to sibling competition (Tangili et al., 2025). In our study population, females have a shorter lifespan than males due to higher actuarial senescence (Briga et al., 2017), and we therefore anticipated the DMSi to be lower in females. However, independent of age, the DMSi in females was only marginally lower compared to males (Table 1a; Fig.4b). Because birds from large broods achieved a shorter lifespan (Briga et al., 2017), we speculated that the DMSi would decline with age when being old would imply having a phenotype that more resembles having grown up in a large brood. However, no significant effects of brood size and age on DMSi were detected (Fig. 4c,d), and this did not change when treating age as a numerical variable instead of a category. Additionally, neither sex nor interactions between age cohort, sex, or brood size significantly influenced DMSi values (Table 1a), suggesting that the effect of brood size on DMSi is consistent across ages and sexes.

**Figure 4.**
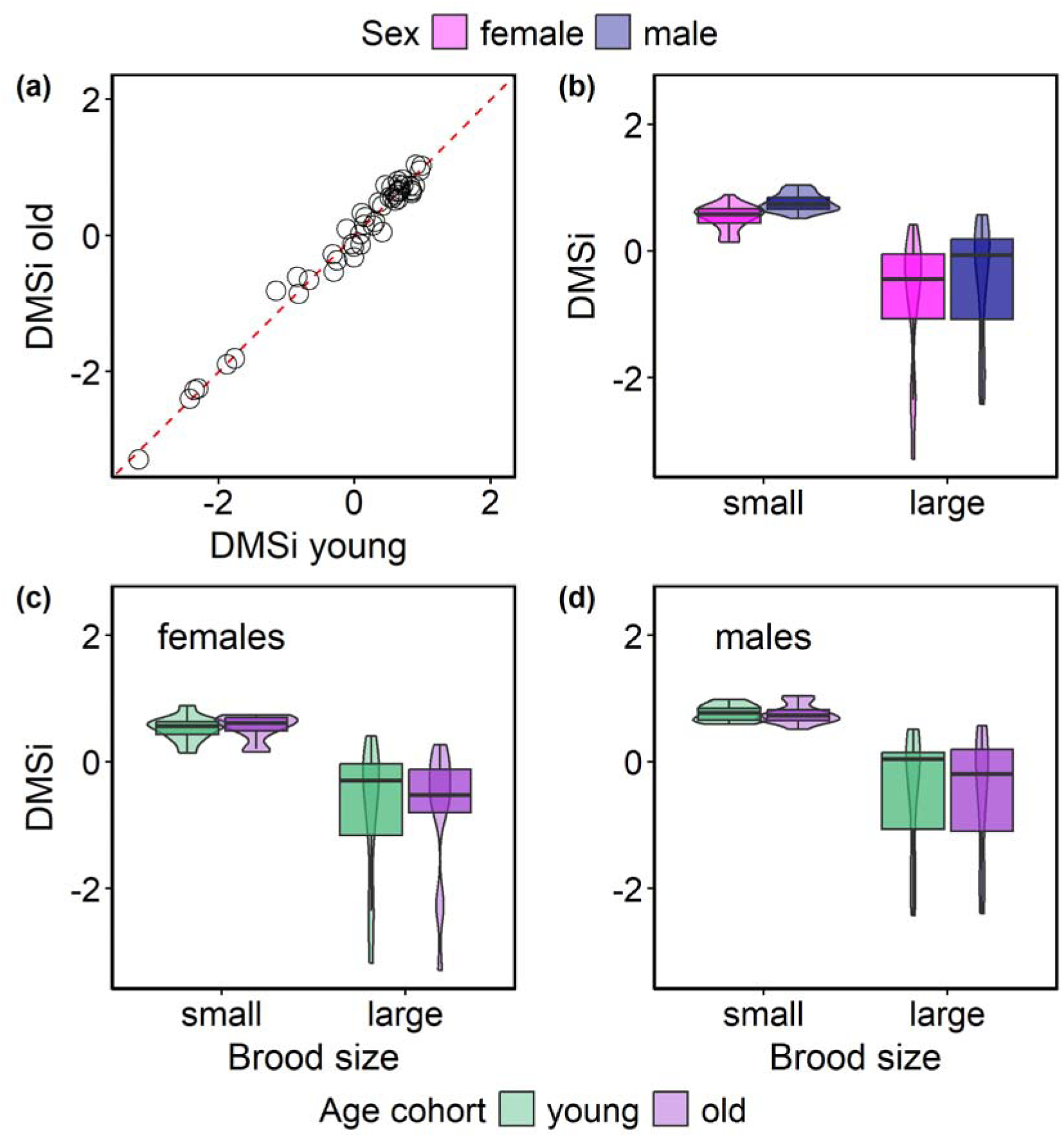
(a) Within individual changes in DMSi (z-scores, r = 0.99, p < 0.001). The dashed line indicates equal values (y=x), so no within-individual change in DMSi with age. Levels of DMSi (z-scores) in zebra finches raised in small and large broods divided by sex in the merged samples (b), and divided by age cohort in females (c) and males (d).

**Table 1.** The DMSi (a) and the SMLmi (b) in relation to brood size and sex. Variables were mean centered with females, small broods and young individuals coded as -0.5 and males, large broods and old individuals coded as +0.5. Bird identity was included as a random effect.

| **(a) DMSi** |  |  |  |  |
| --- | --- | --- | --- | --- |
| **Predictors** | *Estimates (±SE)* | *df* | *t-value* | *Pr(>\|t\|)* |
| Intercept | -0.004(±0.1) | 46 | -0.33 | 0.97 |
| brood size | -1.31(±0.21) | 46 | -6.12 | <0.001 |
| age cohort | -0.005(±0.02) | 46 | -0.24 | 0.81 |
| sex | 0.28(±0.21) | 46 | 1.29 | 0.21 |
| brood size* age cohort | -0.02(±0.04) | 46 | -0.36 | 0.72 |
| brood size* sex | 0.1(±0.43) | 46 | 0.23 | 0.82 |
| age cohort * sex | -0.03(±0.04) | 46 | -0.63 | 0.53 |
| brood size* age cohort *sex | 0.001(±0.09) | 46 | 0.01 | 0.99 |
| **Random Effects** |  |  |  |  |
| BirdID | 0.56 |  |  |  |
| residual | 0.12 |  |  |  |
| **(b) SMLmi** |  |  |  |  |
| **Predictors** | *Estimates (±SE)* | *df* | *t-value* | *Pr(>\|t\|)* |
| Intercept | 6.13e-04 (±0.019) | 47 | 0.03 | 0.98 |
| brood size | 0.295 (±0.038) | 47 | 7.72 | <0.001 |
| age cohort | 0.017 (±0.018) | 47 | 0.94 | 0.35 |
| sex | -0.032 (±0.038) | 47 | -0.83 | 0.41 |
| brood size* age cohort | 0.011 (±0.037) | 47 | 0.32 | 0.76 |
| sex * age cohort | 1.34e-03 (±0.037) | 47 | 0.04 | 0.97 |
| **Random Effects** |  |  |  |  |
| BirdID | 0.01 |  |  |  |
| residual | 0.008 |  |  |  |

### 2.4 Case study: Supervised Machine Learning methylation index (SMLmi)

When predicting brood size using the merged samples per individual, a ridge regression (α=0, using the methylation of all CpG sites to predict brood size with optimized weights) yielded the best result. However, the predictive accuracy was only 42%, i.e. slightly worse than random assignment. This indicates that no individual CpG or small subset of CpGs in the pooled samples was strongly predictive of rearing brood size, which is in line with the identification of DMSs in this data set (Fig. 1). A predictive modeling approach on all individual samples using elastic net regression with leave-one-out cross-validation yielded a different result, predicting rearing brood size with an accuracy of 89% (Fig.5). When inspecting the 11 wrongly assigned samples, it became apparent that eight of these samples were from four individuals of which both samples were wrongly assigned (Table S1). The elastic net model SMLmi utilized the methylation of 455 CpG sites (0.04% of all CpG sites that passed filtering, Fig.S8) spread over 30 autosomal chromosomes and the Z chromosome (Fig.S8b), where it appears that there were more included CpG sites than expected by chance on chromosome 3, and fewer than expected by chance on 1 (Fig.S8a). Five SMLmi CpG sites were shared with the DMSs per sample and three with the male-specific DMSs (Fig.S4). As expected, the SMLmi differed strongly between birds reared in small and large broods but the difference in the SMLmi variance between birds reared in small (0.020) and large (0.023) broods did not approach significance (F = 0.86, 95% C.I. 0.49 - 1.52, d.f. = 49, *p* = 0.61), in contrast to the finding with the DMSi. Age, either as category (young / old) or as continuous variable, and sex did not explain significant variation in the SMLmi (Table 1b).

**Figure 5.**
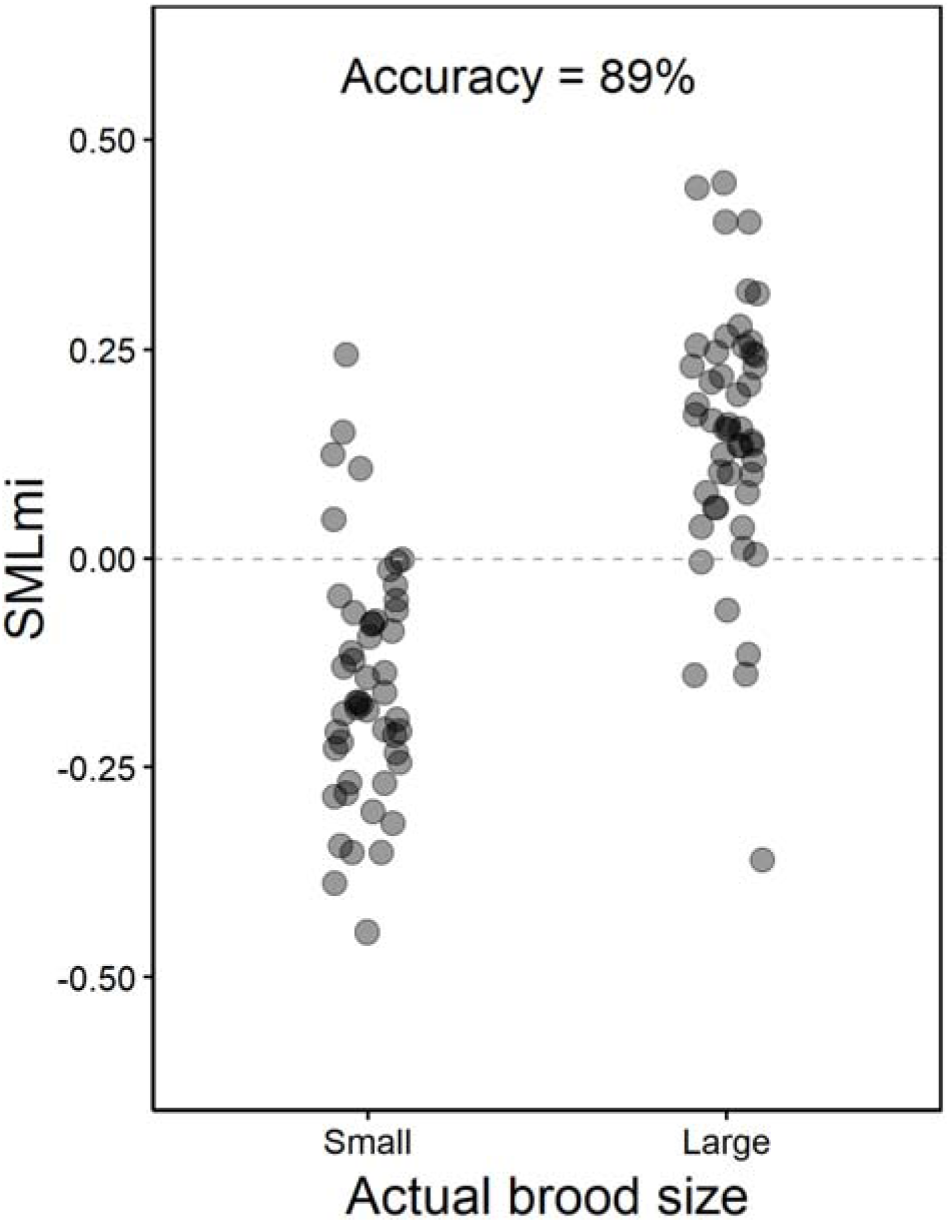
Predicted brood size (small brood: -0.5, large brood: 0.5) from the DNA methylation of 455 CpG sites selected by the final model trained on all samples using the optimized hyperparameters α and λ. The model has a predictive accuracy of 89% (89/100 samples correctly assigned to their brood size). The brood size of the points above the dashed line on the left and under the dashed line on the right was wrongly assigned.

### 2.5 Case study: Comparing indexes

Across samples, we identified a strong negative correlation between DMSi and SMLmi (Fig.6, r = - 0.73, *p* < 0.001). This pattern suggests that both indices capture biologically meaningful epigenetic information about the early-life environment. There was no support for a non-linear relationship, as the inclusion of SMLmi squared or otherwise transformed did not improve the model. When examining this relationship within brood size treatments, the negative association was apparent in both groups (small broods: r = -0.5, *p* < 0.001, large broods: r = -0.56, *p* < 0.001). A z-test revealed that the difference in slopes between small and large broods was statistically significant (z=3.7, *p* < 0.001) suggesting that the relationship between DMSi and SMLmi differed depending on brood size. This finding is in agreement with the larger variance in DMSi among individuals reared in large broods. However, a mixed-effects model including an interaction between SMLmi and brood size did not detect a significant difference in the strength of association between brood sizes (brood size * SMLmi: β = 0.19 ± 0.27, *t* = 0.7, *p* = 0.49), which we attribute to the different position of the two data sets on the X-axis (Fig.6).

**Figure 6.**
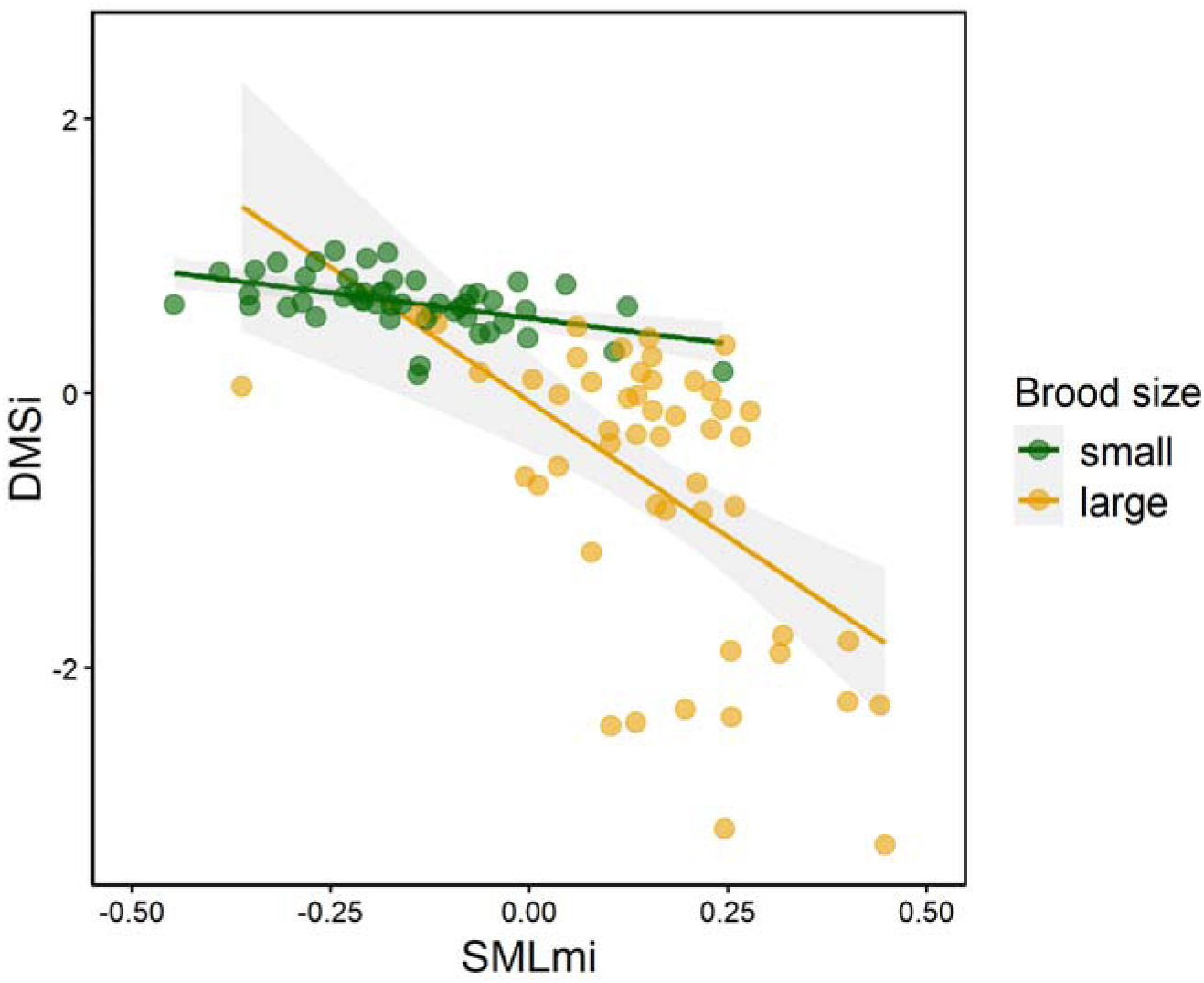
Correlation between DMSi and predicted brood size values (r = -0.73, p < 0.001). For individuals raised in small broods r=-0.05 (p < 0.001) and for individuals raised in large broods r=- 0.56 (p < 0.001). Colors denote the brood size of rearing. Solid line represents the best-fit regression through the data and the colored area is 95% CI.

**Figure 7.**
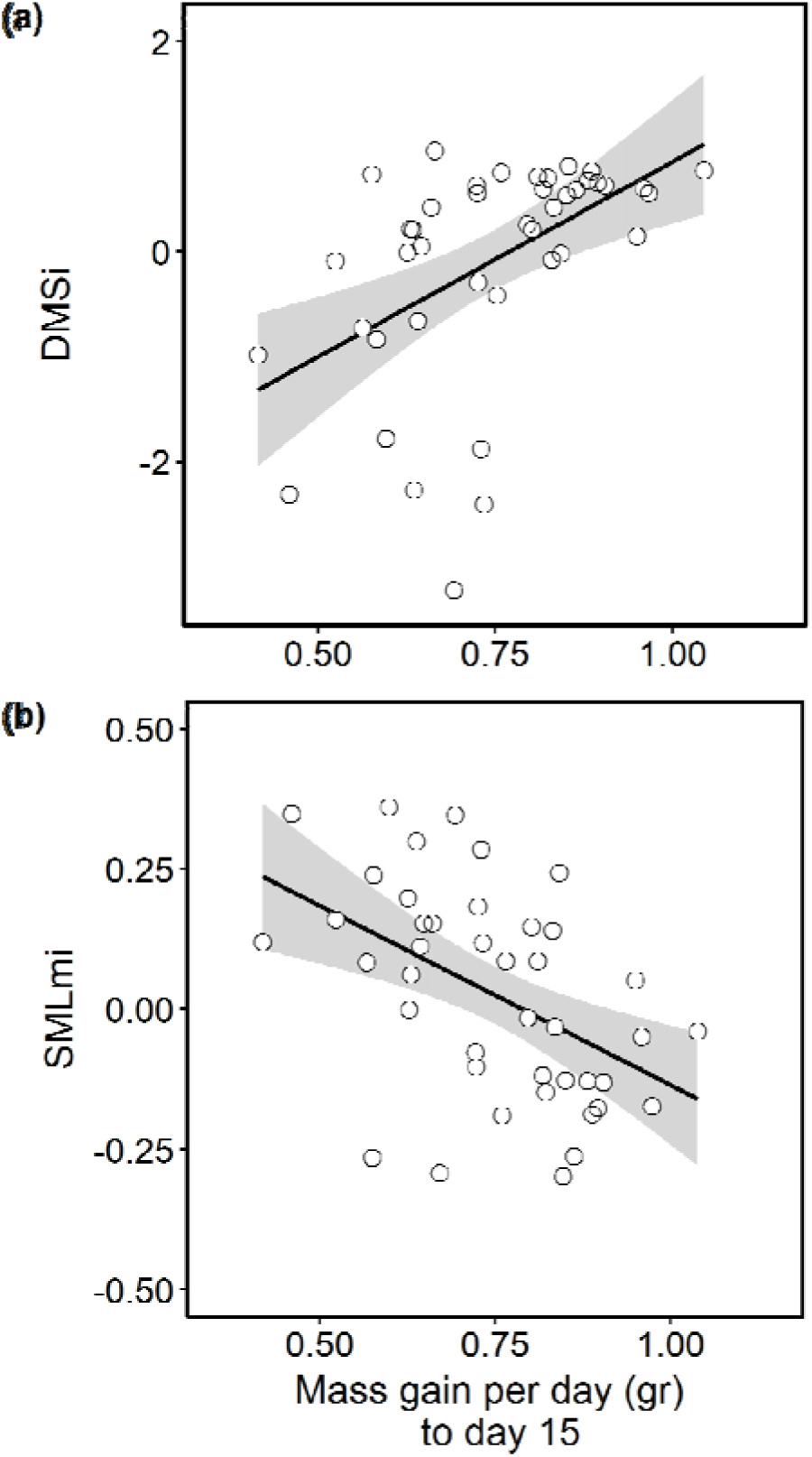
Correlations of DMSi (a) and SMLmi (b) with mass gain per day (gr) from day five to day 15 (N=42). Solid line represents the best-fit regression through the data and the colored area is 95% CI. Growth rate data were corrected for age at measurement using the estimates in Tables S2 and S3.

For most female-specific DMSs the methylation difference was negative and for male-specific DMSs the difference was positive (Fig.S10), meaning that birds in large broods tended to have lower methylation at female specific DMSs while the opposite pattern occurred in male-specific DMSs. The CpGs selected for the SMLmi showed intermediate methylation differences compared to the DMSs (Fig.S10). This distinction reflects the different goals of the two analyses; identifying the CpG sites with the largest methylation differences versus accurately predicting brood size. The CpGs selected to increase SMLmi performance do not necessarily represent the largest differences in methylation between treatments. This is to be expected, with machine learning algorithms selecting CpG sites that add the largest amount of information given the other selected CpG sites. CpG sites where DNAm responds strongly to treatment (i.e. brood size in our study) in most cases add relatively little information because DNAm on these sites will be strongly correlated with DNAm on selected sites where DNAm responded to treatment.

### 2.6 Case study: Indexes and early-life growth rate

To assess to what extent the two methylation-based indexes relate to individual variation in early-life conditions on a phenotypic level instead of brood size treatment, we correlated DMSi and SMLmi to pre-fledging growth rates. Early-life growth rate is affected by brood size (Briga et. al 2017) and a robust predictor of fitness prospects across a wide range of species (Douhard et al., 2014; Drake et al., 2025), including birds (Brown et al., 2022). Both indices significantly predicted individual variation in growth rate (Fig.8; Tables S2,3). The partial correlation coefficients for mass gain per day were 0.5 for the DMSi and -0.48 for the SMLmi, and comparison of the AIC values of the predictive models revealed that the DMSi was a slightly better predictor of early-life growth rate than SMLmi (AIC = -47.7 and -46.8, respectively). However, when adding rearing brood size to the models in tables S2 and S3, this caused the index effects to be no longer significant (DMSi: β = 0.02 ± 0.02, *t* = 1.05, *p* = 0.3; SMLmi: β = 0.03 ± 0.05, *t* = 0.53, *p* = 0.6). Thus, both indexes can be predictive of early-life growth, but are not an improvement over knowing the actual early-life conditions, indicating also that growth does not fully characterize the phenotypic consequences of developmental conditions.

**Figure 8.**
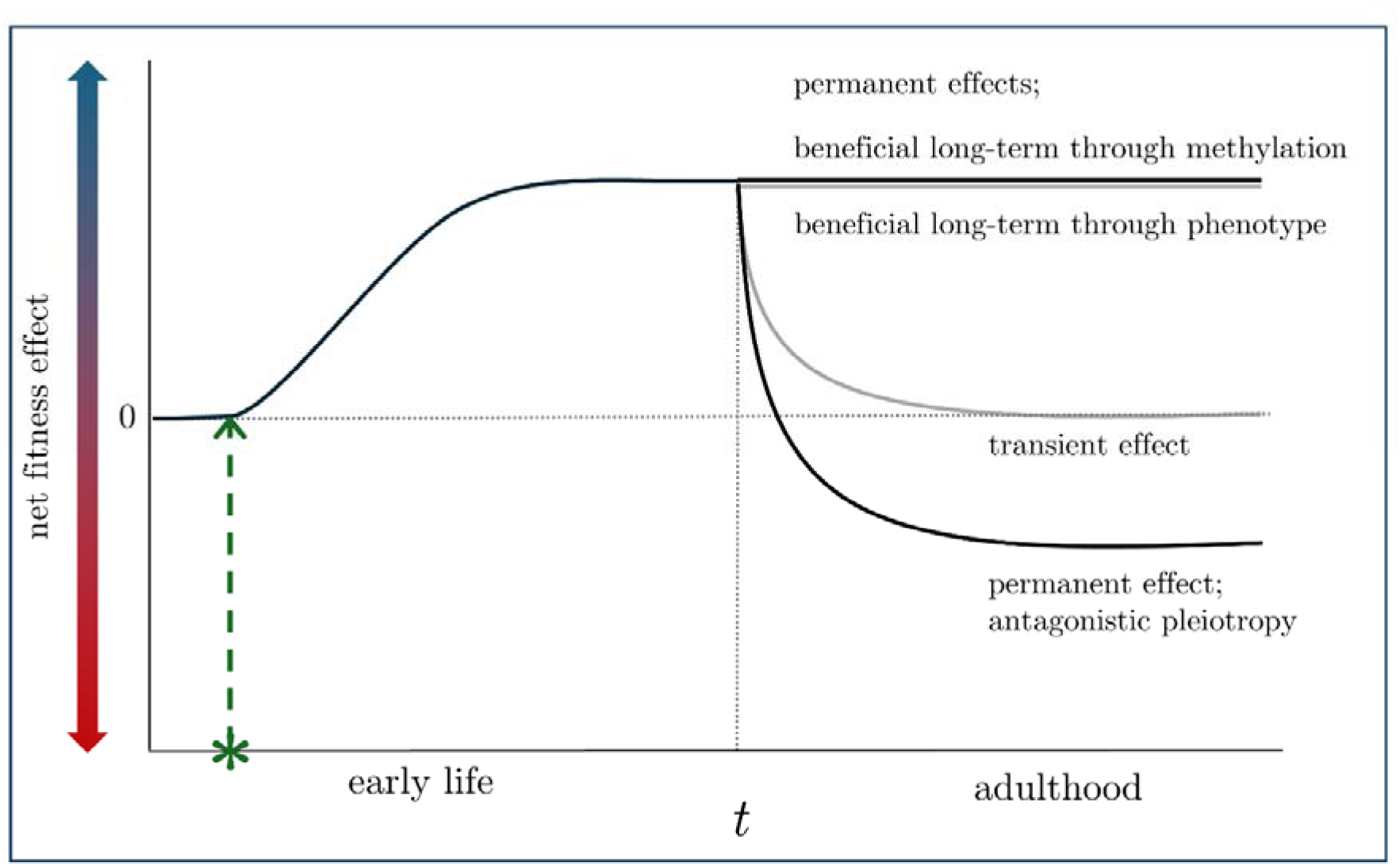
Short and long-term effects of early-life environmental change on fitness through methylation. Solid lines show the net fitness effects of methylation changes caused either by environmental change (green star and vertical arrow) or the onset of adulthood (vertical dotted line). Black solid lines represent the presence and grey solid lines the absence of methylation effects on fitness. For simplicity, the effects of methylation change are here considered either fully permanent or fully transient and the net fitness changes are only due to changes in methylation.

## 3. Discussion

Epigenetic information can be utilized in different ways to understand how environmental factors modulate phenotypic traits in the short and long-term, affecting organismal sensitivity, resilience and adaptability. Studying DNAm at the level of each individual CpG site offers the highest resolution for understanding how DNAm directly affects phenotypes depending on its location, because the methylation of different parts of the genome has different effects on gene expression (Moore et al., 2013). However, even CpG site-level analyses can differ in resolution. Depending on the approach employed to quantify methylation, and the resulting resolution of the data, studies examined methylation in CpG sites across the entire genome, focus on CpG islands, or target specific genomic regions or functionally relevant loci. However, focusing exclusively on genes or genomic regions predicted to be affected by a treatment introduces significant limitations. Novel effects outside of the predefined regions may be overlooked, leading to a possibly misleading understanding of the treatment’s effects on the epigenome. Furthermore, this approach limits the opportunity to use the data in ways not envisaged when designing the study. Taking into consideration all the above, we consider comprehensive, high resolution, genome-wide analyses essential to capture the full spectrum of epigenetic differences between phenotypes that can be leveraged to get insights into the mechanistic underpinnings of phenotypic differences between individuals.

Leveraging whole-genome methylation data from adult zebra finches raised in either small or large broods, we presented a case study to compare different ways to quantify the long-term consequences of early-life developmental conditions on the methylome using longitudinal samples. When identifying DMSs, longitudinal samples, taken at different times from an individual, can either be merged, increasing coverage per site and therefore potentially precision, or analyzed separately. We have here presented analyses on merged samples, separately for the two samples from each individual, taken early and late in life in each individual and separately for males and females.

Judging by the number of DMSs identified, using individual samples was the more powerful approach, and we consider these findings the best estimate of the (long-term) effects of rearing brood size on CpG site specific DNAm. Merging the samples concealed the interesting finding that DNAm differences due to rearing brood size were larger later in life, raising the question whether phenotypic differences due to rearing brood size were also larger later in life. We therefore conclude (not surprisingly), that whether it is recommended to merge longitudinal samples depends on the details of the question that is being addressed, with merging being useful to increase precision when estimating long-term effects, while addressing age-related variation requires samples taken at different times to be analyzed separately.

Statistically, higher coverage at a given CpG site reduces stochastic noise and improves reliability of the methylation estimate but systematic biological differences between the two samples may be averaged out, potentially masking temporal or condition-dependent changes. Whether longitudinal samples were merged or not also affected the development of indexes of early-life conditions. For example, we observed a strong reduction in power of the SMLmi to predict brood size from the merged samples compared to a similar analysis based on the individual samples. This finding indicates that using aggregated longitudinal data might not be suitable for trait prediction and offers further support to the idea that the difference in methylation between individuals raised in small versus large broods varies across different life stages as previously demonstrated in great tits (*Parus major*, Sepers et al., 2023, 2024). We therefore tend towards recommending the calculation of indexes using individual samples, but acknowledge that more comparisons between indexes from merged versus single samples will be necessary before general conclusions can be reached with confidence.

When studying natural populations, developmental conditions of a proportion of individuals will usually not be known, and this proportion can be very high. Our findings suggest that the indexes we calculated based on the effects of manipulated brood size on DNAm offer the opportunity to obtain information on the developmental conditions of individuals first encountered in adulthood. The high predictive accuracy of brood size by SMLmi is encouraging in this respect, as was the strong association between the indexes and early-life growth rate. The actual brood size of rearing was still a better predictor of growth rate than either index, but when direct information about developmental conditions is absent, indexes as developed in this study are possibly the best available alternative, besides traits that can be measured on adults (e.g. morphological traits that are fixed early in life). We recognize, however, that such indexes probably have to be developed and validated for each species or population, and also within populations it remains to be investigated as how stable such indexes are in space and time. Moreover, it remains to be demonstrated that our approach yields similar results in the wild, where environmental heterogeneity is likely to be larger, and survival selection on early life development is stronger. Nonetheless, our findings suggest that at least within the context of an ongoing study where developmental conditions are known for part of the population, DNAm based indexes potentially provide a unique tool to assess the developmental conditions of individuals first encountered as adults.

Indexes integrating DNAm information from many CpG sites are likely to have greater explanatory power with respect to phenotypic variation, when compared to individual CpG sites. Nevertheless, we acknowledge that in order to fully understand the mechanisms through which DNAm mediates phenotypic variation, it is ultimately necessary to investigate the functional roles of individual CpG sites. The SMLmi may then be a better starting point than the DMSi, as that CpG selection method includes validation on independent samples (through the leave-one-out protocol), but for either indexes it would be advisable to test its repeatability using an independent but fully comparable data set. While both indexes we calculated (DMSi & SMLmi) were highly correlated with each other, the two indexes draw information from sets of CpG sites that showed limited overlap (Fig.S9). Instead of considering this a weak point, we see it as broadening of the opportunity to explore the mechanisms underlying phenotypic variation. This divergence generates the possibility that the two indexes capture multiple and possibly partially independent epigenetic signatures that can contribute to the same phenotypic outcomes. Thus, we see the development of DNAm based indexes of phenotypic traits as a stepping stone towards the identification of (networks of) CpG sites that together play an important role in shaping phenotypic variation.

As illustrated by our findings when comparing results based on either single samples or merged longitudinal samples per individual, DNAm is a dynamic and modifiable process, not only during early life and development (Watson et al., 2019) but throughout an individual’s entire lifespan. Information on the dynamics of DNAm at individual CpG sites is limited, in particular in non-model systems, but whether DNAm responses to environmental changes are transient or permanent has large consequences for the evolution of such responses (Fig.8). Environmental changes that induce a phenotypic response through DNAm can impact fitness in different ways, both before and during adulthood, depending on whether the change in DNAm is transient or persistent. In either case, it can be assumed that an environmentally-induced change in methylation provides fitness benefits in the short term when it is an evolved response. However, different scenarios can be envisaged for long-term consequences (Fig.8). In one scenario, the induced DNAm causes a permanent (favorable) change in the phenotype, in which case the fitness benefits are permanent, regardless of whether the change in DNAm is maintained, making this scenario less relevant in the context of this discussion. Alternatively, methylation patterns can be stably maintained when the different DNAm state is also beneficial in adulthood, maintaining also associated fitness benefits. When the effect on DNAm is transient, and no longer beneficial, the DNAm change may diminish as the individual matures, becoming neutral with respect to fitness. Lastly, an antagonistic pleiotropic effect can arise, where due to a constraint on phenotypic flexibility a change in DNAm that is beneficial early in life is permanent and becomes deleterious in later life. The above scenarios showcase how early-life methylation modifications resulting from early-life environmental exposures can be reinforced, attenuated or altered by later-life exposures, which can lead to complex, lifelong phenotypic effects. A better understanding of the extent to which DNAm dynamics are transient or permanent will be essential for our understanding of the impact of DNAm on the relationship between environment, aging, phenotype and fitness.

### 3.1 Limitations

While present findings reveal a clear association between site-specific DNAm and brood size of rearing, the mechanistic pathways through which early-life conditions affect DNAm remain unclear. In avian species, a harsh development can have profound, direct effects on Darwinian fitness, influencing traits such as growth (de Kogel, 1997; Nilsson & Svensson, 2001), recruitment (Schwagmeyer & Mock, 2008), nestling survival (Mock et al., 2009), foraging behavior (Andrews et al., 2015) and (lifetime) reproductive success (Eastwood et al., 2019; Spagopoulou et al., 2020). More specifically, in zebra finches being raised in a large has been found to affect nestling behavior (Tangili et al., 2025), social integration (Gerritsma et al., 2025), mate preference (Holveck & Riebel, 2010) and lifespan (Briga et al., 2017). Future research efforts should aim to disentangle whether these behavioral and physiological consequences arise through direct effects of the development on the methylome or that they reflect secondary pathways mediated by altered physiological and social dynamics. A relevant concern in this context is that DNAm effects of developmental conditions may arise indirectly, in response to phenotypic changes triggered by the early-life environment (Höglund et al., 2020), instead of underlying the observed phenotypic changes.

The extent to which our findings can be generalized remains an open question, given that the identification of differential methylation can be highly study and/or method-dependent, in that for example data type, analysis tools, filtering choices, statistical framework, and significance thresholds can strongly influence the results. For example, in great tits, initial approaches using pooled data and relatively underpowered methods suggested significant effects of brood size on nestling DNAm (Sepers et al., 2021), but a subsequent study using more refined methods did not find such an effect (Sepers et al., 2023).

Although our study in effect controlled for genetic background by randomly cross-fostering chicks from different genetic backgrounds to treatments, differences in methylation identified in this study should be interpreted with the caveat that some may reflect underlying genetic differences rather than causal effects of the brood size manipulation. Lastly, the modest sample size prevented subgroup analyses, such as age comparisons separately for the two sexes.

### 3.2 Conclusions

This study demonstrates effects of (manipulated) brood size in which an individual bird is raised on site-specific DNAm in adulthood in a sex-specific manner. Moreover, we have here illustrated how methodological choices can have profound effects on the identification of differential methylation, underscoring the need to align analytical pipelines to the specific study aims. It also suggests there is a need for simulation studies to investigate power and biases inherent in the different approaches to the analysis of DNAm. Lastly, the present study highlights the need to understand the causal pathways by which early-life environments affect the phenotype through methylation, including when the methylation marks are established, how stable they are across tissues and life stages, and under which conditions they translate into lasting physiological and behavioral changes.

## Acknowledgements

We thank the Center for Information Technology at the University of Groningen for support and access to Hábrók high-performance computing cluster. We express our gratitude to Ellis Mulder for her help in the laboratory, to Michael Briga and Blanca Jimeno for running the long-term experiment and to the animal caretakers at the University of Groningen as well as numerous students whose invaluable help made this project possible.

## Ethics

All methods and experiments detailed in this manuscript were performed under the approval of the Central Committee for Animal Experiments (Centrale Commissie Dierproeven) of the Netherlands, under license AVD1050020174344.

## Data accessibility

Raw reads of the whole methylomes of all samples are available at the National Center for Biotechnology Information under BioProject ID PRJNA1108628. The bioinformatics code is available in https://datadryad.org/dataset/doi:10.5061/dryad.wm37pvmw8, the R code and intermediate data files used in this study are available in http://datadryad.org/share/LINK_NOT_FOR_PUBLICATION/ZxtGRgc_KmEIC-S2ZKYyAj0cLw_CdjcNT1UmfT4XeT8 (reviewer URL).

## Author’s contributions

MT and SV designed the study. MT performed the analyses. MT and SV wrote the manuscript with input from all authors. All authors approved the final version of the manuscript.

## Competing interests

We have no competing interests.

## Funding

Contributions by PJP were supported by the European Union’s Horizon 2020 Research and Innovation Programme under the Marie Skłodowska-Curie grant agreement no. 813383, and the University of Groningen.

## Supplementary Information

**Figure S1.**
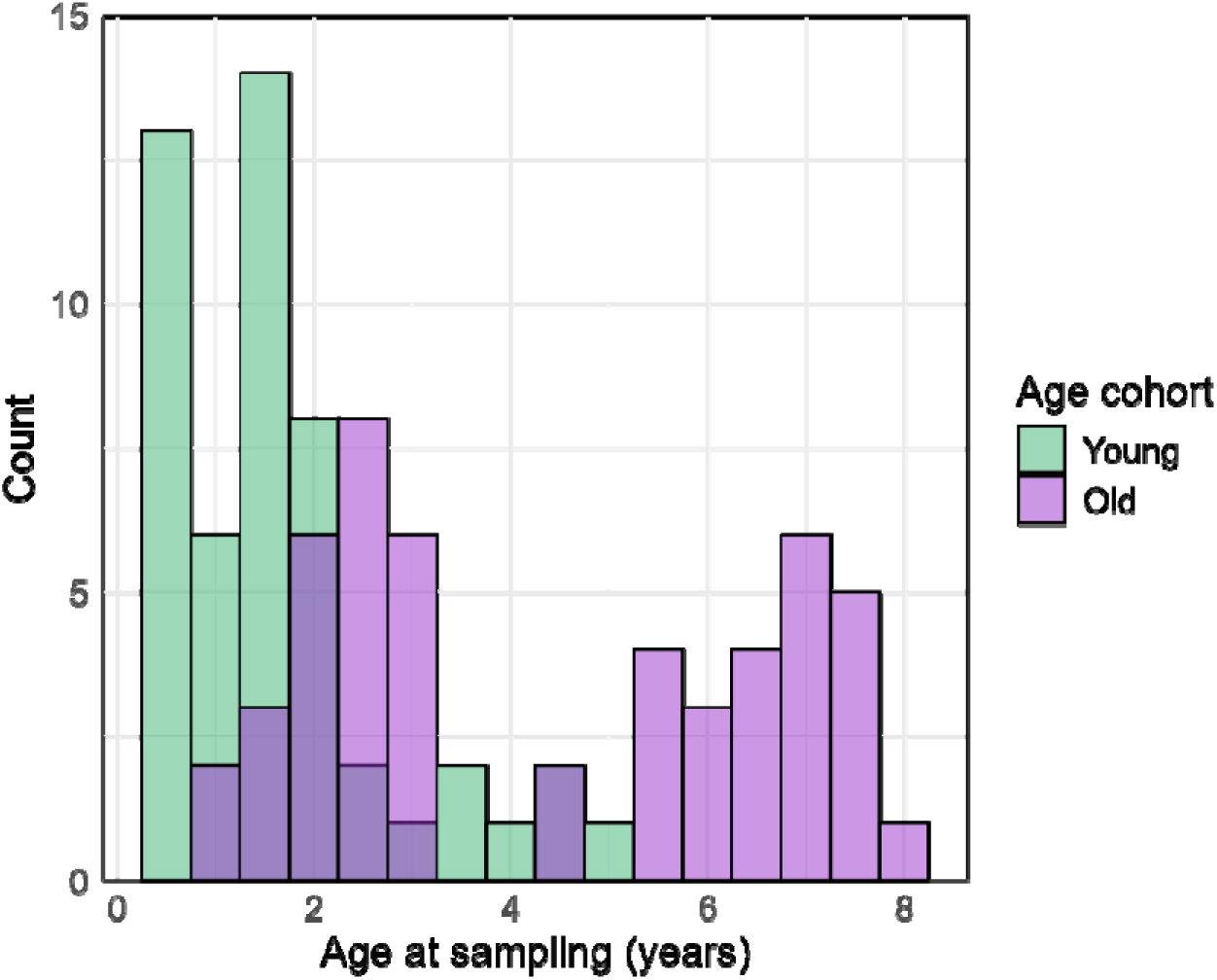
Age at sampling (years) distribution of samples included in the analyses. Color denotes relative time of sampling (young=first sample from an individual, old= second sample of an individual).

**Fig. S2.**
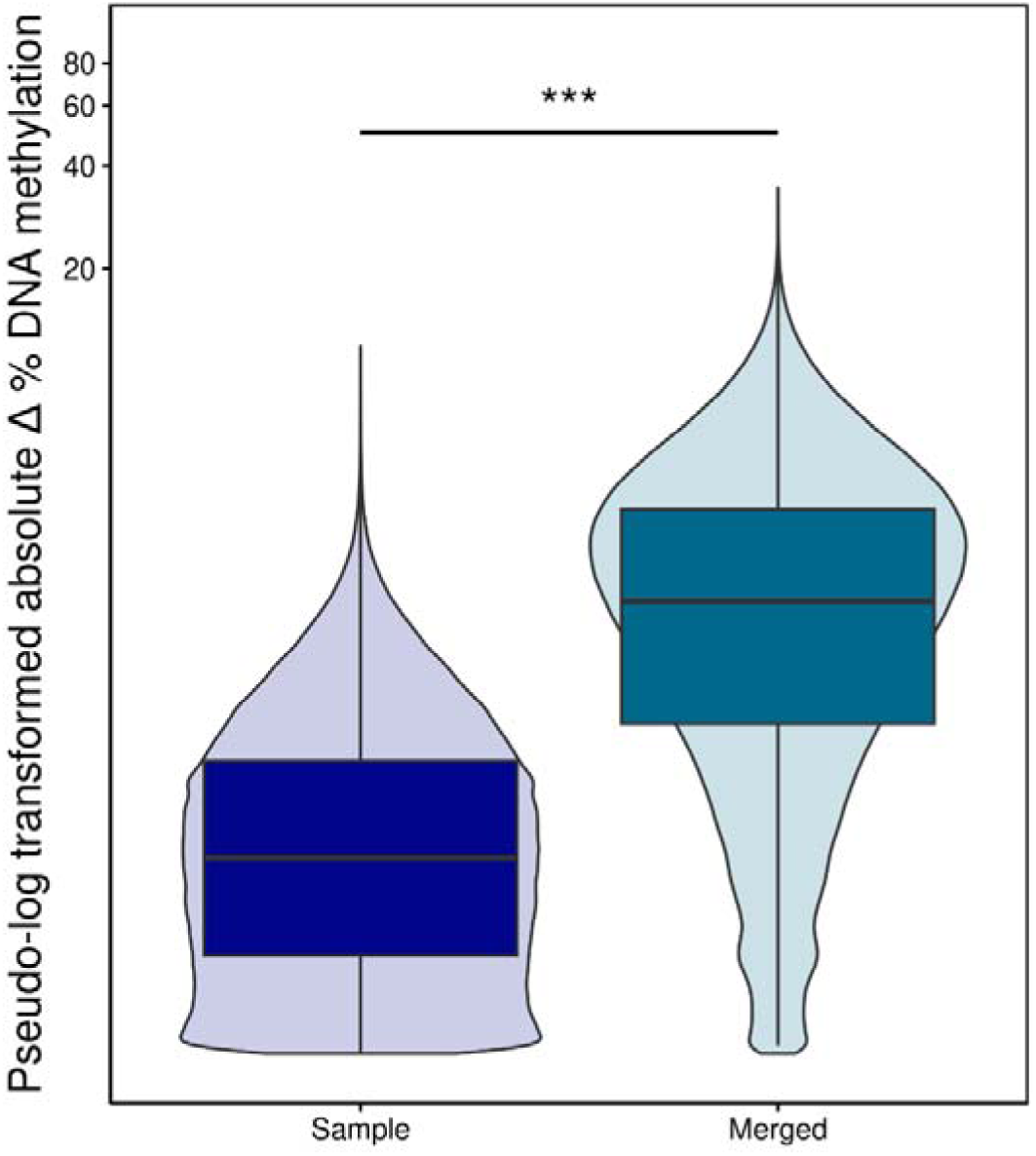
Distribution of CpG-site specific absolute methylation differences between zebra finches raised in small and large broods for using individual samples (N=1,114,704) and for individuals merged together (N=2,085,308). Violin plot widths represent the data density and boxplots show the median and quartiles. The y-axis values were pseudo-log transformed.

**Figure S3.**
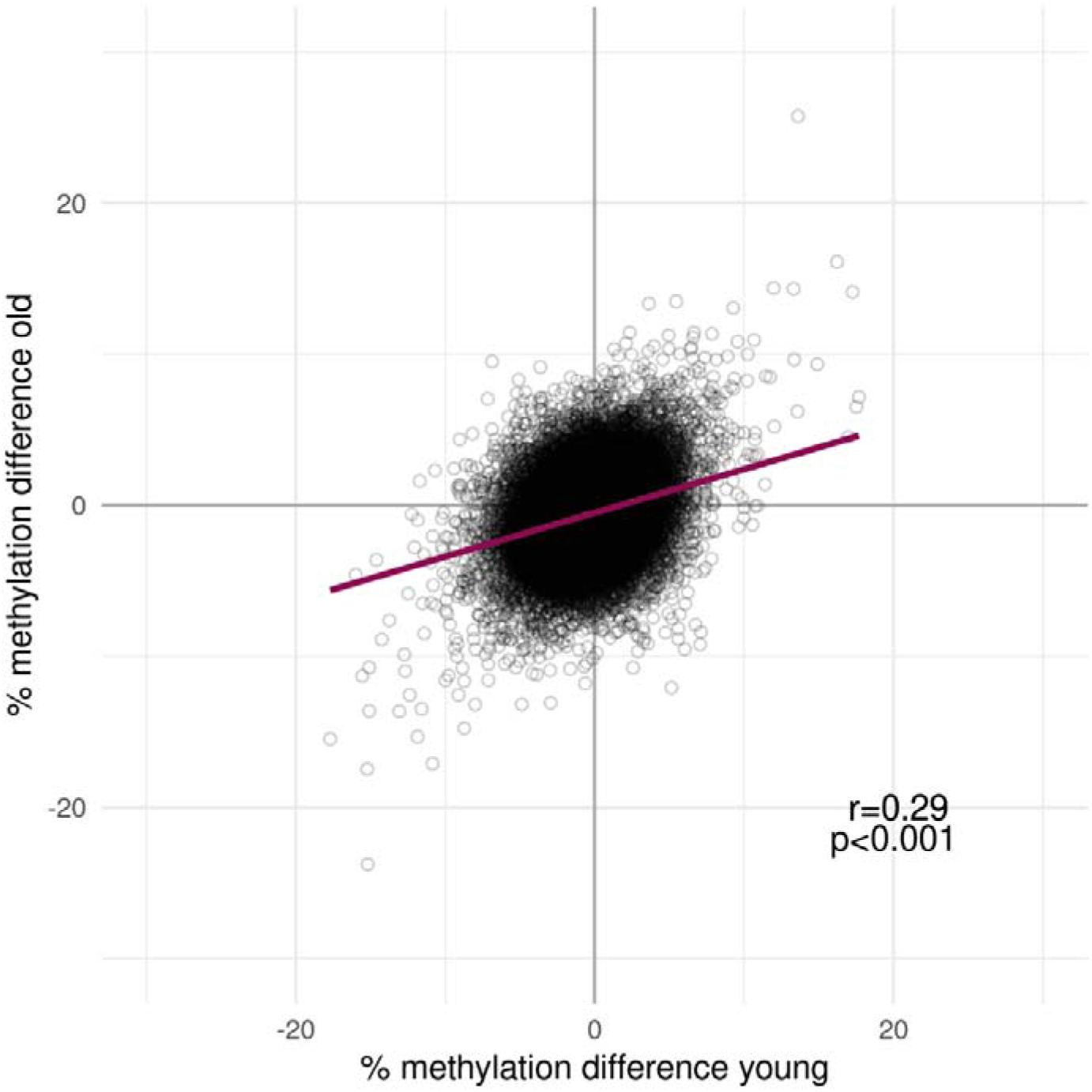
Percent DNA methylation difference between individuals raised in small vs. large broods in old against young individuals. Each data point represents one CpG site. Solid line represents the best-fit regression through the data.

**Figure S4.**
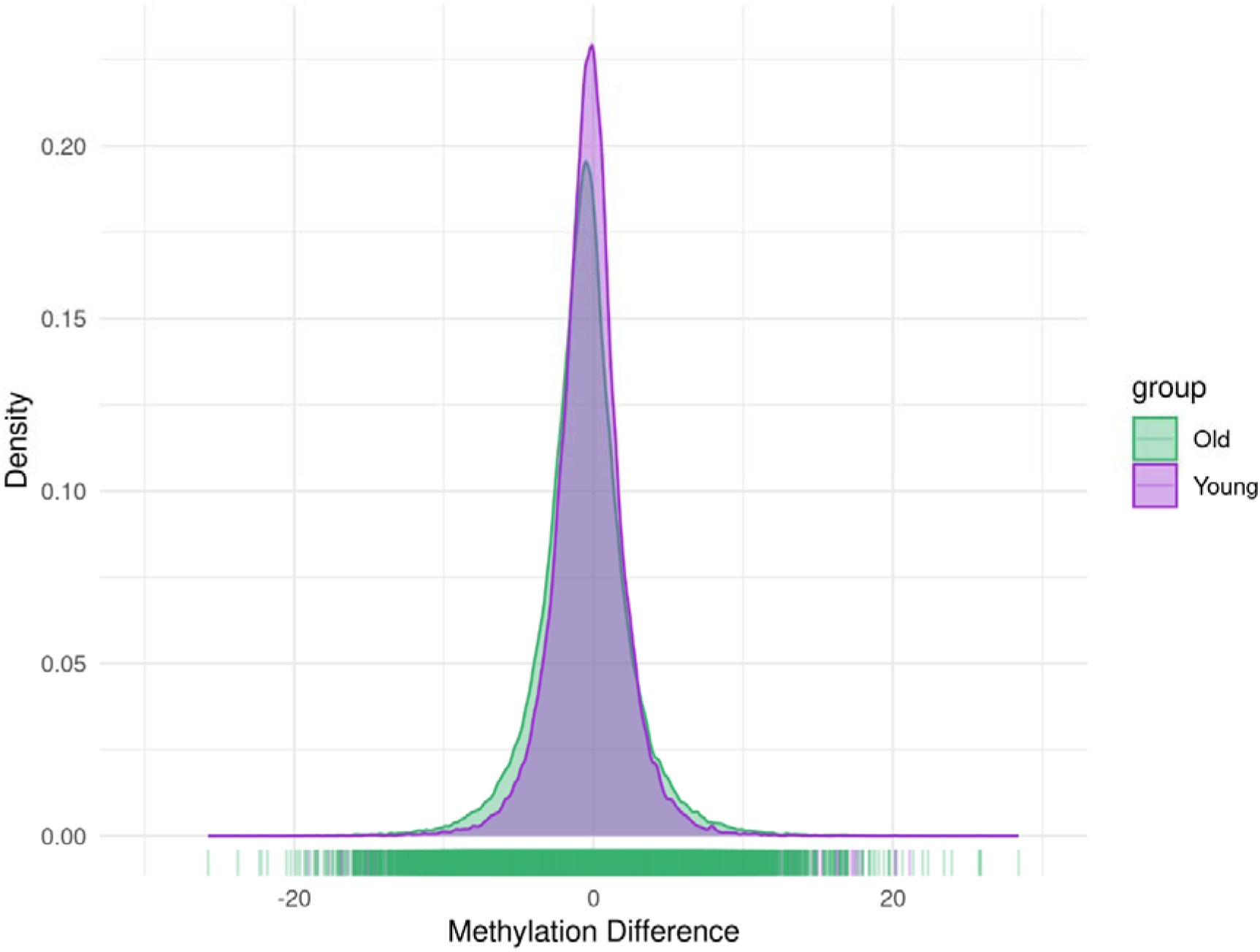
Density plot of percent methylation difference between individuals raised in small vs. large broods in young and old birds.

**Figure S5.**
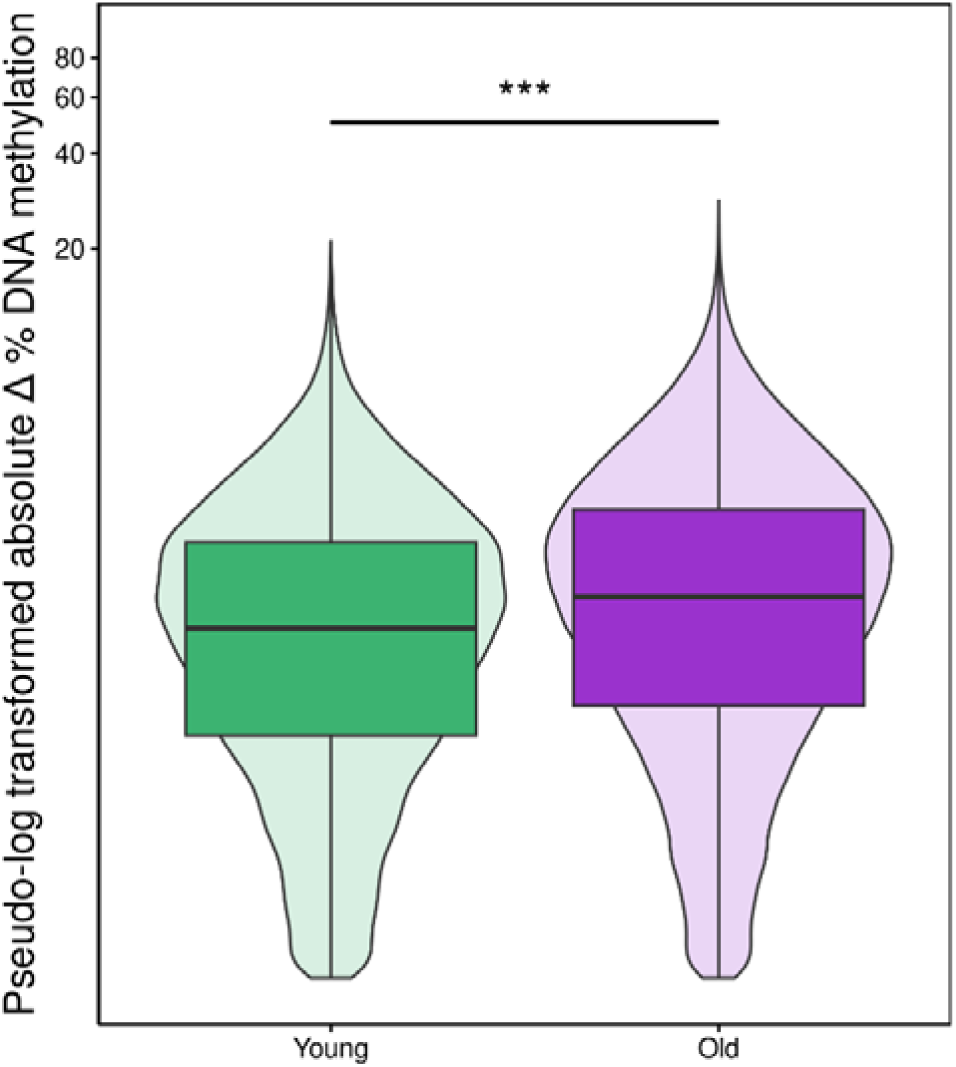
Distribution of CpG-site specific absolute methylation differences between zebra finches raised in small and large broods for the young (N= 60,046) and for the old cohort (N= 82,273). Violin plot widths represent the data density and boxplots show the median and quartiles. The y-axis values were pseudo-log transformed.

**Figure S6.**
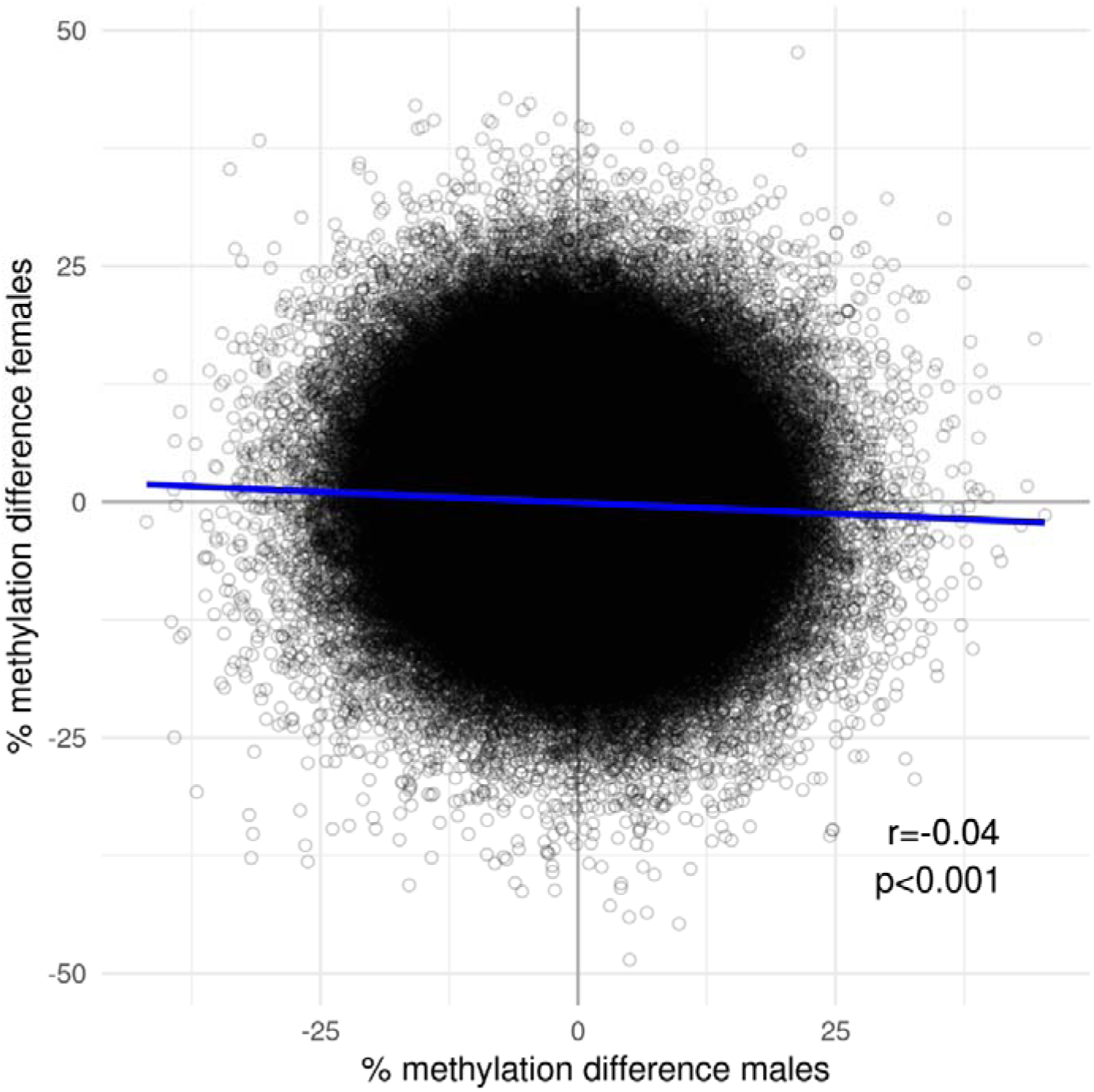
Relationship between CpG methylation differences (N = 1,108,263) observed in male and female zebra finches raised in small and large broods. Solid line represents the best-fit regression through the data.

**Figure S7.**
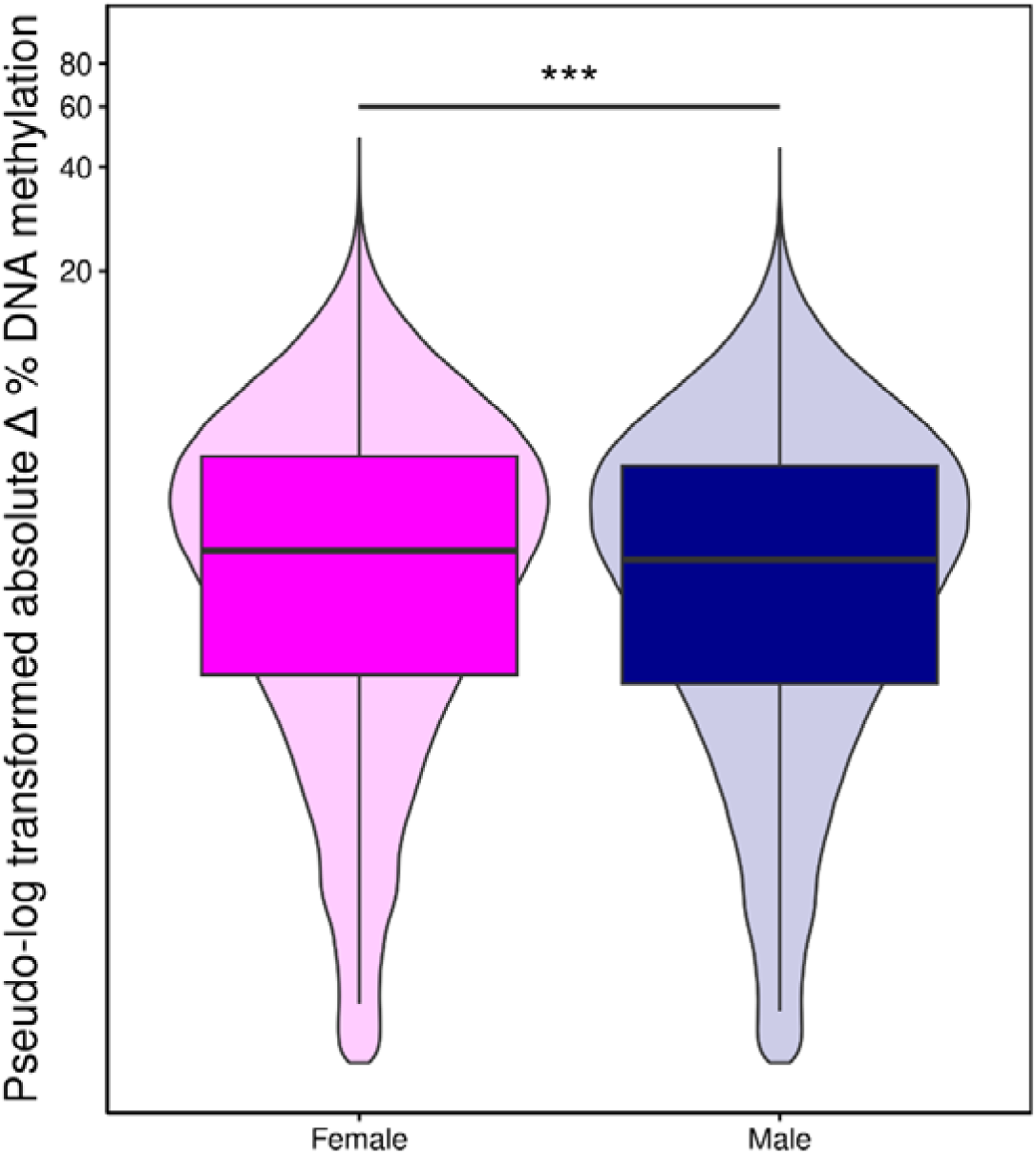
Distribution of CpG-site specific absolute methylation differences between zebra finches raised in small and large broods for females (N= 1,896,932) and only for males (N=1,787,898). Violin plot widths represent the data density and boxplots show the median and quartiles. The y-axis values were pseudo-log transformed.

**Table S1.** Sample information and predicted brood size. Small brood size is coded as -0.5 and large brood size as +0.5. Accuracy denotes whether or not the sample was assigned to the correct brood size. Predictions were considered correct if they matched the direction of the actual brood size deviation (i.e., both positive or both negative).

| Sample | BirdID | Brood size | Predicted brood size | Accuracy |
| --- | --- | --- | --- | --- |
| 1 | 1810 | -0.5 | 0.190 | Incorrect |
| 2 | 1825 | -0.5 | -0.030 | Correct |
| 3 | 1904 | 0.5 | -0.106 | Incorrect |
| 4 | 1907 | -0.5 | -0.046 | Correct |
| 5 | 1920 | 0.5 | 0.068 | Correct |
| 6 | 1944 | -0.5 | -0.114 | Correct |
| 7 | 1953 | -0.5 | -0.085 | Correct |
| 8 | 1983 | 0.5 | 0.103 | Correct |
| 9 | 2045 | -0.5 | 0.003 | Incorrect |
| 10 | 2053 | -0.5 | -0.059 | Correct |
| 11 | 2058 | -0.5 | -0.078 | Correct |
| 12 | 2069 | 0.5 | 0.010 | Correct |
| 13 | 2087 | -0.5 | -0.078 | Correct |
| 14 | 2108 | 0.5 | -0.030 | Incorrect |
| 15 | 2124 | -0.5 | -0.060 | Correct |
| 16 | 2142 | -0.5 | -0.086 | Correct |
| 17 | 2153 | 0.5 | -0.093 | Incorrect |
| 18 | 2155 | -0.5 | -0.190 | Correct |
| 19 | 2163 | 0.5 | 0.141 | Correct |
| 20 | 2175 | -0.5 | -0.055 | Correct |
| 21 | 2229 | 0.5 | 0.086 | Correct |
| 22 | 2260 | 0.5 | 0.125 | Correct |
| 23 | 2291 | 0.5 | 0.084 | Correct |
| 24 | 2236 | 0.5 | 0.195 | Correct |
| 25 | 2259 | 0.5 | 0.086 | Correct |
| 26 | 1983 | 0.5 | 0.028 | Correct |
| 27 | 2429 | 0.5 | 0.111 | Correct |
| 28 | 2473 | -0.5 | -0.342 | Correct |
| 29 | 1923 | 0.5 | 0.115 | Correct |
| 30 | 1942 | 0.5 | 0.001 | Correct |
| 31 | 2058 | -0.5 | -0.175 | Correct |
| 32 | 2269 | -0.5 | -0.078 | Correct |
| 33 | 2087 | -0.5 | -0.179 | Correct |
| 34 | 2172 | -0.5 | -0.079 | Correct |
| 35 | 2288 | -0.5 | -0.028 | Correct |
| 36 | 2423 | 0.5 | 0.268 | Correct |
| 37 | 2487 | -0.5 | -0.024 | Correct |
| 38 | 2045 | -0.5 | -0.108 | Correct |
| 39 | 2257 | -0.5 | -0.049 | Correct |
| 40 | 2277 | 0.5 | -0.060 | Incorrect |
| 41 | 2291 | 0.5 | 0.050 | Correct |
| 42 | 2418 | 0.5 | 0.065 | Correct |
| 43 | 2390 | -0.5 | -0.055 | Correct |
| 44 | 2511 | -0.5 | -0.205 | Correct |
| 45 | 1953 | -0.5 | -0.019 | Correct |
| 46 | 2163 | 0.5 | 0.092 | Correct |
| 47 | 2247 | 0.5 | 0.141 | Correct |
| 48 | 2260 | 0.5 | 0.200 | Correct |
| 49 | 2463 | 0.5 | -0.028 | Incorrect |
| 50 | 1920 | 0.5 | 0.097 | Correct |
| 51 | 2155 | -0.5 | -0.043 | Correct |
| 52 | 1810 | -0.5 | 0.148 | Incorrect |
| 53 | 1825 | -0.5 | -0.060 | Correct |
| 54 | 2053 | -0.5 | 0.089 | Incorrect |
| 55 | 2069 | 0.5 | 0.079 | Correct |
| 56 | 2153 | 0.5 | -0.116 | Incorrect |
| 57 | 2508 | 0.5 | 0.024 | Correct |
| 58 | 2508 | 0.5 | 0.102 | Correct |
| 59 | 2116 | 0.5 | -0.003 | Incorrect |
| 60 | 2259 | 0.5 | 0.147 | Correct |
| 61 | 2423 | 0.5 | 0.293 | Correct |
| 62 | 2257 | -0.5 | -0.023 | Correct |
| 63 | 1897 | 0.5 | 0.091 | Correct |
| 64 | 2463 | 0.5 | -0.035 | Incorrect |
| 65 | 2038 | 0.5 | -0.046 | Incorrect |
| 66 | 2229 | 0.5 | 0.044 | Correct |
| 67 | 2277 | 0.5 | 0.055 | Correct |
| 68 | 2354 | -0.5 | -0.127 | Correct |
| 69 | 2124 | -0.5 | -0.109 | Correct |
| 70 | 1907 | -0.5 | 0.012 | Incorrect |
| 71 | 2511 | -0.5 | -0.296 | Correct |
| 72 | 2429 | 0.5 | 0.280 | Correct |
| 73 | 2473 | -0.5 | -0.081 | Correct |
| 74 | 2035 | 0.5 | 0.069 | Correct |
| 75 | 1944 | -0.5 | 0.071 | Incorrect |
| 76 | 1897 | 0.5 | 0.084 | Correct |
| 77 | 2108 | 0.5 | 0.014 | Correct |
| 78 | 1923 | 0.5 | 0.166 | Correct |
| 79 | 2172 | -0.5 | -0.057 | Correct |
| 80 | 2415 | -0.5 | -0.141 | Correct |
| 81 | 2392 | -0.5 | -0.195 | Correct |
| 82 | 2038 | 0.5 | 0.144 | Correct |
| 83 | 1904 | 0.5 | -0.096 | Incorrect |
| 84 | 1942 | 0.5 | 0.094 | Correct |
| 85 | 2470 | -0.5 | -0.120 | Correct |
| 86 | 2142 | -0.5 | -0.044 | Correct |
| 87 | 2487 | -0.5 | 0.034 | Incorrect |
| 88 | 2035 | 0.5 | 0.110 | Correct |
| 89 | 2116 | 0.5 | 0.129 | Correct |
| 90 | 2175 | -0.5 | -0.148 | Correct |
| 91 | 2470 | -0.5 | -0.235 | Correct |
| 92 | 2247 | 0.5 | 0.076 | Correct |
| 93 | 2415 | -0.5 | 0.047 | Incorrect |
| 94 | 2269 | -0.5 | -0.045 | Correct |
| 95 | 2390 | -0.5 | -0.184 | Correct |
| 96 | 2288 | -0.5 | -0.042 | Correct |
| 97 | 2236 | 0.5 | 0.192 | Correct |
| 98 | 2354 | -0.5 | -0.138 | Correct |
| 99 | 2418 | 0.5 | 0.097 | Correct |
| 100 | 2392 | -0.5 | -0.263 | Correct |

**Figure S8.**
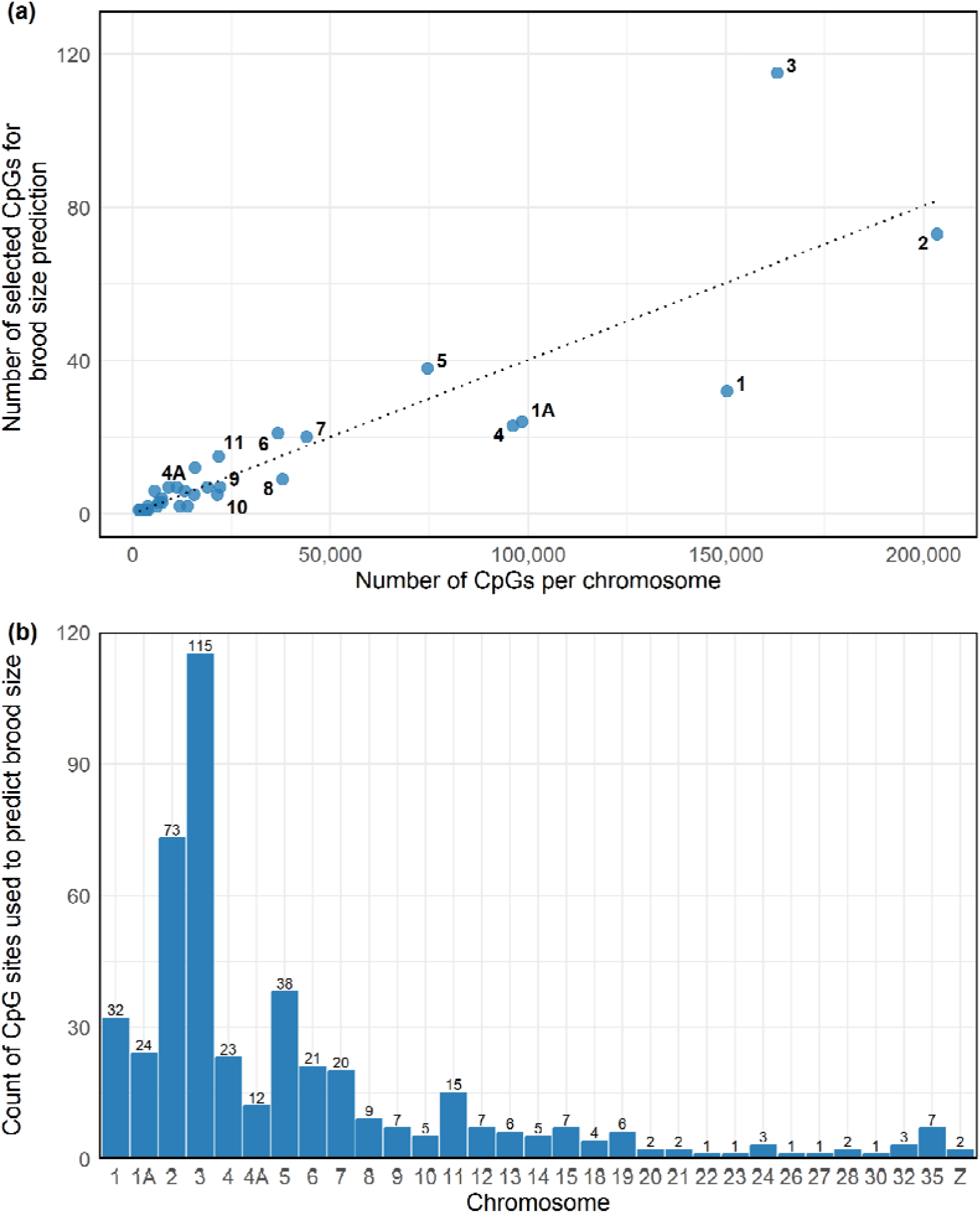
(a) CpG sites in the final brood size prediction model (SMLmi) against total CpG sites per chromosome. The dotted line represents the expected number of CpG sites per chromosome if the CpGs included in the final brood size prediction model were distributed proportionally to the total number of CpG sites in each chromosome. (b) CpG sites in the final brood size prediction model (SMLmi) per chromosome.

**Figure S9.**
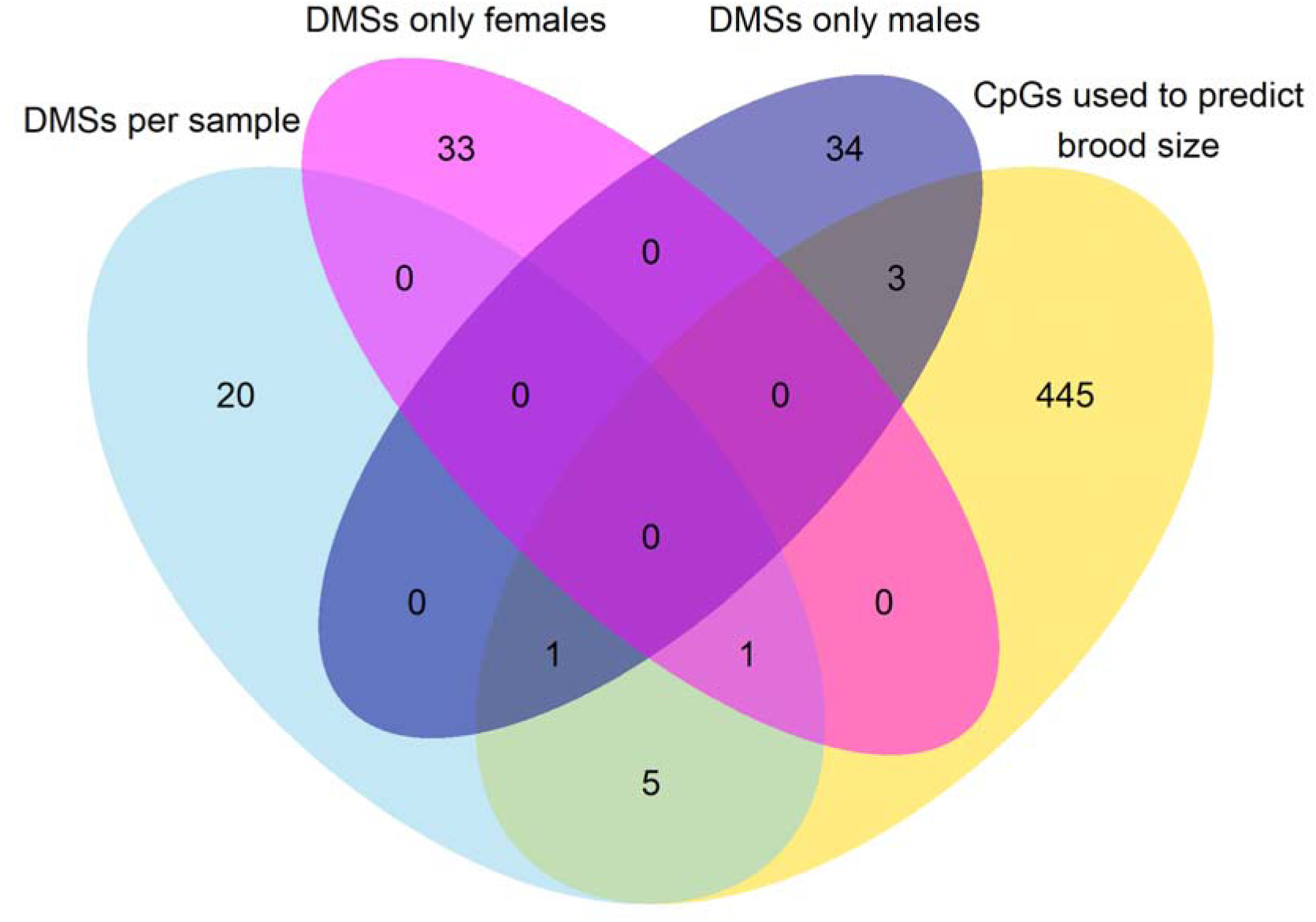
Shared CpG sites between DMSs in the analysis per sample (light blue), in the analysis only for females (pink), in the analysis only for males (blue), and the CpG sites used to predict brood size (i.e. included in the SMLmi, yellow).

**Figure S10.**
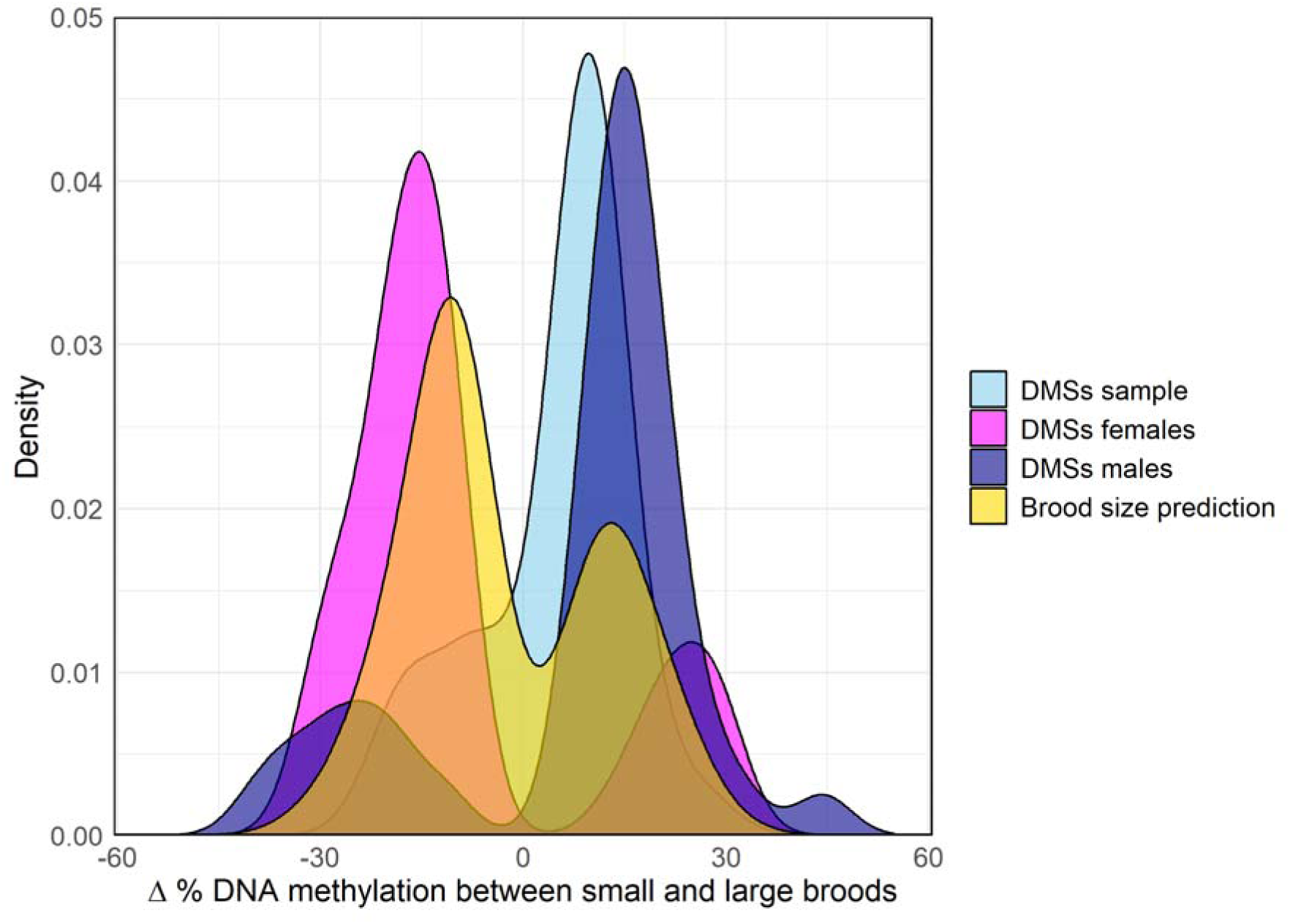
Density plots of methylation differences between individuals raised in small and large broods. Light blue plot: DMSs identified in the analysis per sample (N=27). Pink plot: DMSs identified in the analysis only for females (N=34). Blue plot: DMSs identified in the analysis only for males (N=38). Yellow plot: CpGs selected for the SMLmi (N=455).

**Table S2.**
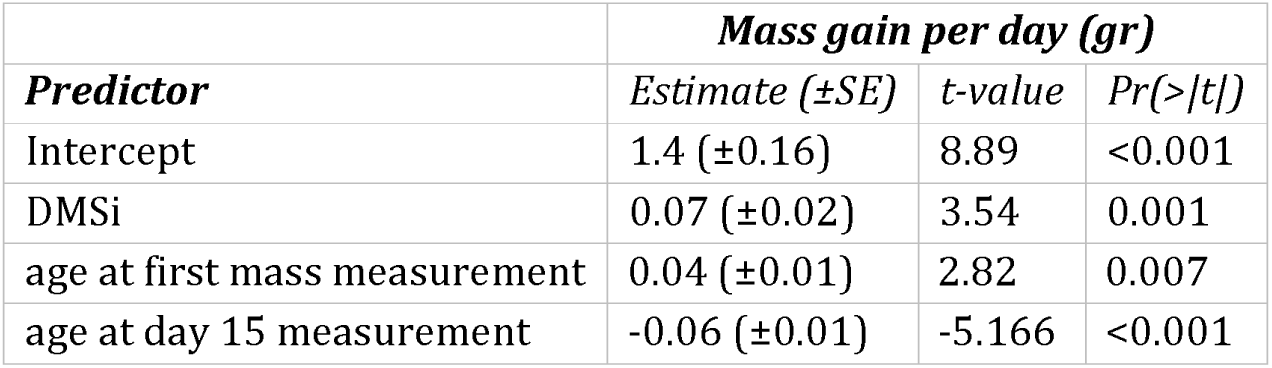
Results of linear model exploring the effects of DMSi and age at measurement on mass gain per day.

**Table S3.** Results of linear model exploring the effects of SMLmi and age at measurement on mass gain per day.

|  | **Mass gain per day (gr)** |  |  |
| --- | --- | --- | --- |
| **Predictor** | *Estimate (±SE)* | *t-value* | *Pr(>\|t\|)* |
| Intercept | 1.43 (±0.16) | 8.82 | <0.001 |
| SMLmi | -0.34 (±0.11) | -3.09 | 0.004 |
| age at first mass measurement | 0.04 (±0.02) | 2.63 | 0.01 |
| age at day 15 measurement | -0.06 (±0.01) | -5.11 | <0.001 |

## Notes

### Competing Interest Statement

The authors have declared no competing interest.

### Summary of Updates

The analysis for the identification of DMSs as well as the development of the DMSi were updated

